# Global ecological and biogeochemical impacts of pelagic tunicates

**DOI:** 10.1101/2022.03.01.482560

**Authors:** Jessica Y. Luo, Charles A. Stock, Natasha Henschke, John P. Dunne, Todd D. O’Brien

## Abstract

The pelagic tunicates, gelatinous zooplankton that include salps, doliolids, and appendicularians, are filter feeding grazers thought to produce a significant amount of particulate organic carbon (POC) detritus. However, traditional sampling methods (i.e., nets), have historically underestimated their abundance, yielding an overall underappreciation of their global biomass and contribution to ocean biogeochemical cycles relative to crustacean zooplankton. As climate change is projected to decrease the average plankton size and POC export from traditional plankton food webs, the ecological and biogeochemical role of pelagic tunicates may increase; yet, pelagic tunicates were not resolved in the previous generation of global earth system climate projections. Here we present a global ocean study using a coupled physical-biogeochemical model to assess the impact of pelagic tunicates in the pelagic food web and biogeochemical cycling. We added two tunicate groups, a large salp/doliolid and a small appendicularian to the NOAA-GFDL Carbon, Ocean Biogeochemistry, and Lower Trophics version 2 (COBALTv2) model, which was originally formulated to represent carbon flows to crustacean zooplankton. The new GZ-COBALT simulation was able to simultaneously satisfy new pelagic tunicate biomass constraints and existing ecosystem constraints, including crustacean zooplankton observations. The model simulated a global tunicate biomass of 0.10 Pg C, annual tunicate production of 0.49 Pg C y^-1^ in the top 100 m, and annual tunicate detritus production of 0.98 Pg C y^-1^ in the top 100 m. Tunicate-mediated export flux was 0.71 Pg C y^-1^, representing 11% of the total export flux past 100 m. Overall export from the euphotic zone remained largely constant, with the GZ-COBALT pe-ratio only increasing 5.3% (from 0.112 to 0.118) compared to the COBALTv2 control. While the bulk of the tunicate-mediated export production resulted from the rerouting of phytoplankton- and mesozooplankton-mediated export, tunicates also shifted the overall balance of the upper oceans away from recycling and towards export. Our results suggest that pelagic tunicates play important trophic roles in both directly competing with microzooplankton and indirectly shunting carbon export away from the microbial loop.

## 1. Introduction

In recent decades, there has been a growing recognition of the prevalence and ecological importance of gelatinous zooplankton (GZ), which include the cnidarian jellyfish, ctenophores, and pelagic tunicates (Hays et al., 2018; Henschke et al., 2016). While they have been a natural component of marine ecosystems extending back to the Cambrian (Hagadorn et al., 2002), their abundance and distributions have been largely overlooked among the public and non-GZ specialists (Condon et al., 2012). This may be due to a combination of the erroneous perception of GZ being “trophic dead-ends” within marine food webs (Hays et al., 2018; Lynam et al., 2006; Verity and Smetacek, 1996), as well as systematic biases in sampling leading to overall under-sampling of their biomass. Net-based sampling has been prevalent for over a century, and is very effective for sampling fish and hard-bodied, crustacean zooplankton (Wiebe and Benfield, 2003). Unfortunately, the fragile gelatinous zooplankton are often broken apart in nets, yielding a *ca.* 3 fold underestimation of their abundance and a *ca.* 10 fold (range: 5-15) underestimation of their carbon biomass relative to non-extractive, optical sampling (Remsen et al., 2004). The rise of in-situ plankton imaging systems have resulted in improved estimates of GZ abundance and distribution (e.g., Luo et al., 2014), yielding advances in understanding of their food-web interactions and biogeochemical impacts (Greer et al., 2021; Robison, 2005; Smith Jr et al., 2014). Combined with the increase in ecosystem-level studies of GZ (e.g., Stukel et al., 2021) and technological advances revealing the importance of GZ as food for higher trophic levels (Hays et al., 2018), there has been an overall paradigm shift in our understanding of the importance and role of gelatinous zooplankton within marine ecosystems.

Amongst zooplankton, GZ are notable for their high clearance rates and boom-and-bust population dynamics, which yield mass mortality events (“jelly-falls”) that sink rapidly through the water column (Acuña et al., 2011; Billett et al., 2006; Lebrato et al., 2012; Lucas and Dawson, 2014). As a result, models have estimated, based on population densities and allometric scaling of ecological and physiological rates, a large contribution (e.g., 1.6-5.2 Pg C y^-1^ in Luo et al. 2020) of GZ-mediated carbon in the global biological pump, with their relative impact increasing with depth (Lebrato et al., 2019; Luo et al., 2020). However, these studies have been done independently of other biogeochemical and ecological constraints (i.e., “offline” calculations), which may yield unrealistic estimates of GZ contributions to marine ecosystems. Indeed, in a new model that includes cnidarian jellyfish as a plankton functional type (PlankTOM11), global export production exhibited very modest increases (+0.1 Pg C y^-1^), suggesting that the online inclusion of a jellyfish class does not by itself substantially increase total export production (Wright et al., 2021). Of the three major groups of GZ considered by Luo et al. (2020), cnidarian jellyfish had the largest standing stock biomass but pelagic tunicates, despite their much lower biomass, had over two times more sinking detritus than cnidarians and ctenophores combined.

These pelagic tunicates, small filter-feeders including appendicularians, salps, doliolids, and pyrosomes, are less conspicuous than the cnidarians and ctenophores, but are highly significant components of marine ecosystems due to their low trophic position, high clearance rates, and fast sinking fecal pellets (Andersen, 1998; Berline et al., 2011; Henschke et al., 2016; Hopcroft and Roff, 1998). Compared to crustacean mesozooplankton such as copepods which feed at predator to prey size ratios ranging from 5:1 to 100:1 (Hansen et al., 1994), pelagic tunicates can feed at predator to prey size ratios ranging from 10:1 to 10000:1 (Conley et al., 2018). Salps pump water through their fine mucous meshes that can filter submicron particles such as bacteria and picoplankton; they are able to sustain the entirety of their energetic demands by grazing on these size classes alone (Sutherland et al., 2010). Using both external and internal filters to feed, appendicularians have the widest range of predator to prey size ratios (exceeding 2500:1 to lower than 10:1), and thus can feed on organisms ranging from 0.2-20 µm in size (Conley et al., 2018; Deibel and Lee, 1992; Fernández et al., 2004). The offline Luo et al. (2020) model estimated that, due to these feeding characteristics, these pelagic tunicates consume between 3.8-8.3 Pg C y^-1^ in prey. Of this consumption, approximately 12-17% were later consumed by higher trophic level predators, and 55-60% became detritus.

Global ocean biogeochemical and marine ecosystem models typically represent marine food webs with roughly linear food chains of phytoplankton to zooplankton to (implicit) fish (Kearney et al., 2021). The traditional inclusion of multiple zooplankton groups has been to distinguish between zooplankton size, such as microzooplankton and mesozooplankton, with the latter parameterized using crustacean zooplankton measurements (Aumont et al., 2015; Buitenhuis et al., 2006; Stock and Dunne, 2010; Ward et al., 2012). Even in models with complex food-webs, predator to prey size ratios beyond ∼50:1 (cf. Hansen et al., 1994) are rarely considered. As such, current ocean biogeochemical models typically represent a marine ecosystem in which crustaceans dominate zooplankton ecology. While this view represents certain ecosystems well (Pershing and Stamieszkin, 2020), globally, there is a tension between the traditional, crustacean-dominated zooplankton view of marine ecosystems and a shifting paradigm that emphasizes the role of GZ. Unfortunately, GZ-focused offline models are unable to reconcile this tension, as evidenced by high GZ-mediated global ingestion and production rates (Luo et al., 2020) that may not support primary and secondary production rates for crustacean zooplankton consistent with observations. Additionally, offline studies are limited in the capacity to explore factors underlying observed GZ niches, and how GZ impacts emergent food web patterns. These challenges are compounded by stubborn limitations in GZ observations which are patchy and inconsistently sampled.

In this study, we added two explicit zooplankton functional types that represent thaliaceans (salps, doliolids, pyrosomes) and appendicularians into Carbon, Ocean Biogeochemistry, and Lower Trophics version 2 (COBALTv2; Stock et al., 2020), a global model designed to represent a “traditional” marine ecosystem dominated by crustacean zooplankton. We ask the following four questions:

1) Can simulations capture the magnitude and gradients of observed GZ biomass across ocean biomes and along productivity gradients, after accounting for approximately an order of magnitude under-sampling by nets?
2) Can simulations reconcile recent evidence for the importance of GZ with established evidence for the prominence of crustacean zooplankton in biogeochemical cycles and the plankton food web?
3) How does a simulation of GZ-modulated export that satisfies multiple food web constraints compare with offline estimates?
4) What is the net impact of GZ zooplankton on the partitioning of carbon flows between recycling, carbon export and energy flows to higher trophic levels?

## 2. Methods

As a brief overview of the methods, we begin with a description of plankton food web dynamics within the original COBALTv2 marine ecosystem model (Stock et al., 2014a, 2020), followed by the GZ additions that comprise GZ-COBALT: small and large pelagic tunicates. We first describe the baseline parameterization of GZ-COBALT, and then a few sensitivity experiments that explore parts of the parameter space and particular elements of the tunicate groups that make them distinct from crustacean zooplankton. Next, we detail the physical framework of the model. Finally, we describe the construction of a validation dataset for the two new GZ groups and the identification of an emergent relationship that contrasts gelatinous against crustacean zooplankton. The model is validated against multiple constraints, comprising new and established ecological and biogeochemical datasets.

### 2.1 COBALT Ecosystem Model

We use the COBALTv2 marine ecosystem model (Stock et al., 2020) as our baseline model configuration, with slight modifications. COBALTv2 is a 33-tracer, intermediate complexity model, representing biogeochemical cycles of carbon, alkalinity, oxygen, nitrogen, phosphorus, iron, silica, calcium carbonate, and lithogenic materials. The food web consists of three phytoplankton and three zooplankton functional types, as well as a free-living heterotrophic bacteria group. Two phytoplankton size classes, a small (smp) and large (lgp) phytoplankton are represented, as well as diazotrophs (diaz), parameterized as a large *Trichodesmium* group. The small phytoplankton type includes cyanobacteria and other phytoplankton, up to 10 µm in equivalent spherical diameter (ESD), and the large phytoplankton type represents diatoms and other large phytoplankton from 10-100 µm in ESD. The different sized phytoplankton are parametrized to capture size-based contrasts in nutrient uptake, light harvesting, carbon to chlorophyll ratios, and susceptibility to microzooplankton grazing (Edwards et al., 2015, 2012; Hansen et al., 1994; Munk and Riley, 1952), such that the small phytoplankton are more successful in the low nutrient, seasonally stable subtropical gyres, and large phytoplankton are more competitive in the highly seasonal, high nutrient oceans (Stock et al., 2014a, 2020).

The base configuration of COBALTv2 contains three zooplankton size classes: a microzooplankton and two size classes of crustacean mesozooplankton. Microzooplankton (smz; < 200 µm ESD) include ciliates and heterotrophic nanoflagellates, medium zooplankton (mdz; 200-2000 µm ESD) are small mesozooplankton and represent small to medium-bodied copepods, and large zooplankton (lgz; 2 – 20 mm ESD) are large mesozooplankton that represent large copepods and krill (Stock et al., 2014a). Predator-prey relationships are also largely based on size, with microzooplankton predating on bacteria and small phytoplankton, small mesozooplankton predating on diazotrophs, large phytoplankton, and microzooplankton, and large mesozooplankton predating on diazotrophs, large phytoplankton, and small mesozooplankton (Fig. 1a). Grazing is modeled as a Hollings Type II function with density-dependent switching (Stock et al., 2008), with maximum biomass specific grazing rates decreasing with increasing zooplankton size (Hansen et al., 1997). Grazing half-saturation constants do not vary between the zooplankton classes, and are tuned to reproduce observed patterns in phytoplankton biomass and turnover rates (Stock and Dunne, 2010).

**Figure 1.**
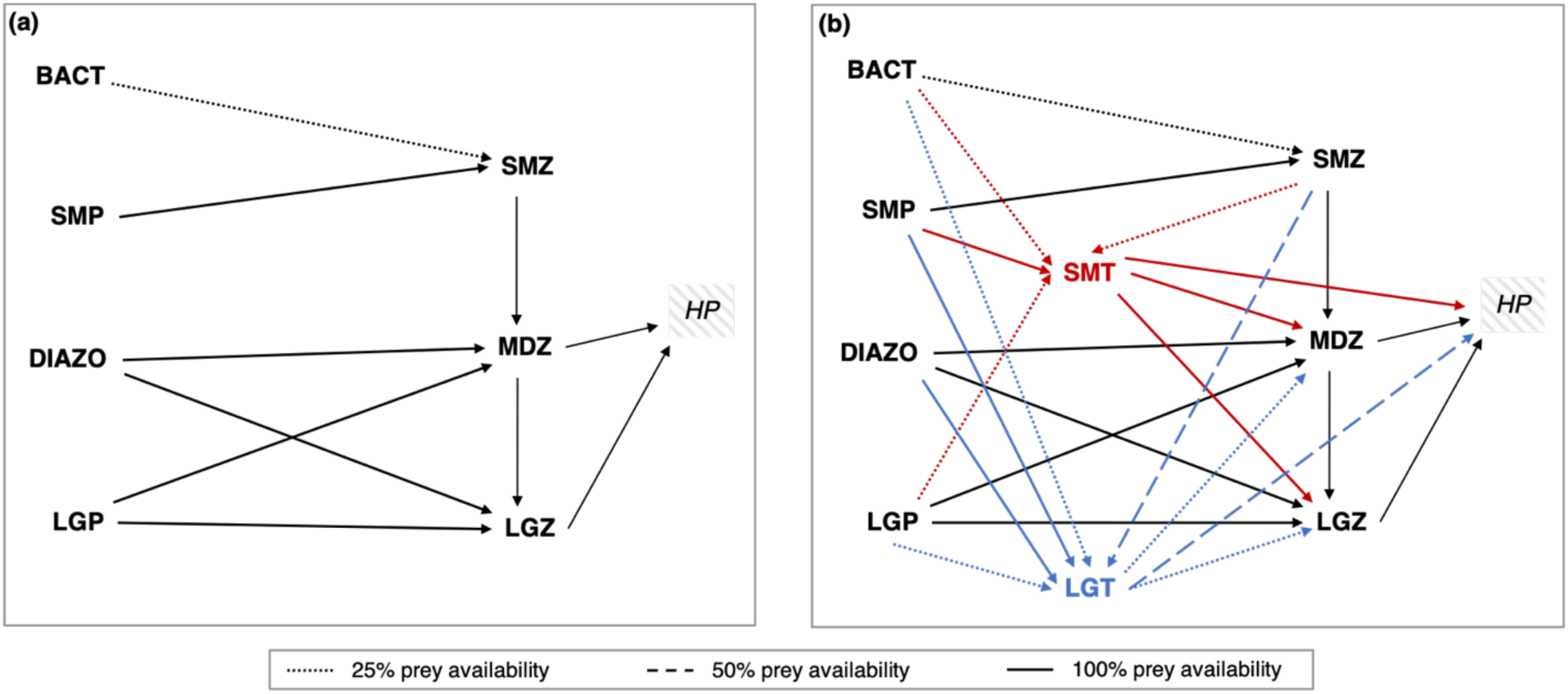
Food web structure of (a) COBALTv2 base model, and (b) GZ-COBALT model. Additional functional groups in GZ-COBALT include a small (0.3-3 mm body length) and large (3-300 mm body length) tunicate, abbreviated as SMT and LGT, respectively. Bact = free living bacteria, SMP = small phytoplankton (< 10 µm ESD), Diazo = diazotrophs (e.g., *Trichodesmium*), LGP = large phytoplankton (10-100 µm ESD), SMZ = small zooplankton (< 200 µm ESD), MDZ = medium zooplankton (200-2000 µm ESD), LGZ = large zooplankton (2-20 mm ESD), and HP = higher trophic-level predators. Note that within typical zooplankton size categories, small zooplankton would be considered microzooplankton, and medium and large zooplankton would be considered mesozooplankton.

Zooplankton grazing was assimilated at an efficiency (AE) of 0.7, with the non-assimilated grazing partitioned into dissolved and particulate (detritus) matter, depending on size. Detritus partitioning of non-assimilated matter were: 1/6 for microzooplankton, 2/3 for medium, and 1 for large zooplankton, and the remainder separated between labile (70%), semi-labile (20%) and semi-refractory (10%) dissolved matter. Assimilated matter was partitioned into respiration (basal and active) and zooplankton production. Basal respiration is proportional to biomass, whereas active respiration is proportional to ingestion rates (see eq. 2, Table 1). When respiration rates exceed assimilated ingestion, production becomes negative and recontributes to zooplankton mortality. The temperature dependence of biological rates (*T_f_*; unitless) including phytoplankton nutrient uptake and growth and zooplankton grazing, is determined by a common Eppley (1972) curve:

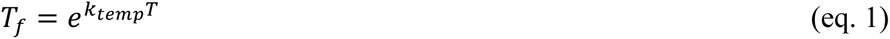

where *T* is the temperature in °C, and *k_temp_* is the temperature scaling factor, set to 0.063 °C^-1^. This corresponds to a Q_10_ of 1.88, representing a near doubling of rates for every 10°C temperature increase. While there is evidence indicating that there may be different temperature dependencies for phytoplankton vs. zooplankton processes (Lopez-Urrutia et al., 2006; Rose and Caron, 2007), as well as for tunicates (Andersen and Nival, 1986; Iguchi and Ikeda, 2004; Lombard et al., 2005), we followed the practice established in COBALTv1 of adopting a single temperature scaling for all plankton types for the sake of parsimony (Stock et al., 2014a).

**Table 1.** GZ-COBALT zooplankton ingestion, respiration, and aggregation parameters. Plankton functional types (PFT)s: smt = small tunicates, lgt = large tunicates, smz = small zooplankton, mdz = medium zooplankton, and lgz = large zooplankton.

| Parameter | Name (Units) | PFT | Values | References |
| --- | --- | --- | --- | --- |
| $I_{max}$ | Maximum ingestion rate at 0°C (day<sup>-1</sup>) | smt | 1.875 | (Lombard et al., 2009a) |
|  |  | lgt | 0.55 | (Luo et al., 2020; Madin and Deibel, 1998) |
|  |  | smz | 1.42 | (Hansen et al., 1997; Stock et al., 2020; Stock and Dunne, 2010) |
|  |  | mdz | 0.57 |  |
|  |  | lgz | 0.23 |  |
| $K_i$ | Ingestion half-saturation constant (mg C m<sup>-3</sup>) | smt, lgt | 250 | (Acuña and Kiefer, 2000; Gibson and Paffenhöfer, 2000) |
|  |  | smz, mdz, lgz | 102 | (Stock et al., 2014a, 2020) |
| $b_{resp}$ | Basal respiration rate at 0°C (day<sup>-1</sup>) | smt | 0.06 | (Lombard et al., 2005) |
|  |  | lgt | 0.008 | (Luo et al., 2020; Madin and Deibel, 1998); tuned to allow for sufficient biomass in subtropical gyres |
|  |  | smz | 0.018 | (Hansen et al., 1997; Stock et al., 2020; Stock and Dunne, 2010) |
|  |  | mdz | 0.008 |  |
|  |  | lgz | 0.0032 |  |
| $f_{aresp}$ | Fraction of ingestion rate for active respiration (unitless) | smt, lgt | 0.15 | See text; also tuned to allow for sufficient biomass in subtropical gyres |
|  |  | smz, mdz, lgz | 0.3 | (Stock et al., 2014a, 2020) |
| $AE_{min}, AE_{max}$ | Minimum and maximum assimilation efficiency (unitless) | smt, lgt | 0.25, 0.8 | (Lombard et al., 2009a; Madin and Deibel, 1998; Pakhomov et al., 2006) |
|  |  | smz, mdz, lgz | 0.7, 0.7 | (Stock et al., 2014a, 2020) |
| $K_{AE}$ | Assimilation efficiency half-saturation constant (mg C m<sup>-3</sup>) | smt | 110 | (Berline et al., 2011; Lombard et al., 2009a) |
|  |  | lgt | 215 | (Harbison et al., 1986) |
|  |  | smz, mdz, lgz | N/A | N/A |
| $f_{agg}$ | Fraction of maximum ingestion at which aggregation mortality is suppressed (unitless) | lgt | 0.1 | See text |
| $m_{agg}$ | Aggregation loss rate (m<sup>3</sup> mg C<sup>-1</sup> d<sup>-1</sup>) | lgt | 1.0E-3 | Calibrated to achieve jelly-falls representing 35% of total mortality |

### 2.2 Gelatinous Zooplankton in COBALT (GZ-COBALT)

We introduced two new pelagic tunicate groups into COBALTv2 (Fig. 1b): small tunicates (smt), which represents Appendicularians and range from 0.3-3 mm in body length, and large tunicates (lgt), which represents Thaliaceans such as salps, doliolids, and small pyrosomes that span 3-300 mm in body length. Appendicularians are small, free-swimming organisms that produce gelatinous houses for filter-feeding, which are discarded when clogged and re-created multiple times per day. Thaliaceans are also filter-feeders, but unlike appendicularians, are colonial (though salps and doliolids have solitary life stages), form rapidly sinking fecal pellets (Perissinotto and Pakhomov, 1998), and exhibit mass die-offs (jelly-falls; Henschke et al., 2013).

The generalized equation that describes the dynamics of the modeled gelatinous zooplankton (*GZ*; mg C m^-3^) is as follows:

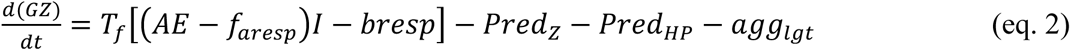

where *T_f_* is the effect of temperature on biological rates (eq. 1; unitless), represented via the Eppley function (see Section 2.1), *AE* is the assimilation efficiency (eq. 6; unitless), *f_aresp_* is the fraction of ingestion that goes to active respiration (unitless), *I* is ingestion (mg C m^-3^ d^-1^), and *bresp* is the basal respiration rate (day^-1^). Population losses include the loss to other zooplankton predators (*Pred_Z_*; mg C m^-3^ d^-1^), the loss to higher trophic level predators (*Pred_HP_*; mg C m^-3^ d^-1^), and aggregation mortality in large tunicates (*agg_lgt_*; mg C m^-3^ d^-1^; eqs. 7-8) which represent jelly-falls.

As described in the previous COBALT documentation papers (Stock et al., 2014a, 2020), zooplankton ingestion is modeled as a Hollings type II saturating functional response dependent on its maximum ingestion rate (*I_max_*; day^-1^), half-saturation constant (*K_i_*; mg C m^-3^), and total prey resources (*P*; mg C m^-3^):

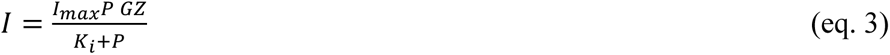

where *P* is represented as the sum of all individual prey types, *p_i_*, multiplied by Φ_i_ (unitless, range from 0-1), which is the prey availability with a switching factor (Stock et al., 2008):

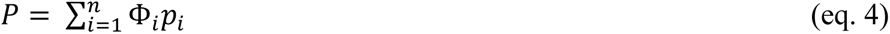

The prey availability for different predator-prey pairs are described in the text in Section 2.2.1. As the modeled GZ are both predators and prey of different plankton types, they also experience predation losses from other zooplankton (*Pred_z_*) and higher trophic level predators (*Pred_HP_*). The formulation of the aggregation losses (*agg_lgt_*) representing jelly-falls are discussed in Section 2.2.4. This loss term is the only new loss term in GZ-COBALT relative to COBALTv2.

Additionally, compared to the COBALTv2 formulation (Stock et al., 2020), here we have separated out the maximum gross growth efficiency (*gge_max_*) into its component parts, which is the assimilation efficiency (*AE*) and the fraction of ingestion to active respiration (*f_aresp_*). This is to enable a dynamic AE that varies with prey concentration. These parameterizations are further discussed in Sections 2.2.2 and 2.2.3.

#### 2.2.1 GZ food web structure

The pelagic tunicates were inserted into the COBALTv2 model as summarized in Fig. 1b. Where possible we will also indicate the short abbreviation for the modeled plankton group in parentheses to avoid confusion caused by multiple equivalent names. Pelagic tunicates have the widest prey-to-predator size ratios among zooplankton (Conley et al., 2018), with appendicularians predating on organisms ranging from 0.04 – 20% of its size, approximately 0.2 – 20 µm in length (Deibel and Lee, 1992; Fernández et al., 2004; Lombard et al., 2011). Small tunicates (smt) in the baseline model setting are thus able to consume bacteria (bact), small phytoplankton (smp), large phytoplankton (lgp), and microzooplankton (smz). This is consistent with previous efforts that included appendicularians in a NPZD-type model (Berline et al., 2011) wherein small tunicates consumed small phytoplankton, which lie near the center of their prey kernel, with high preference (100% prey availability), and all others with low preference (25% prey availability). Appendicularians are important prey for many invertebrates and fish, often contributing large proportions of the diets of medusae jellyfish and many larval fish species (Gorsky and Fenaux, 1998; Llopiz et al., 2010; Purcell et al., 2005). Thus, small tunicates were predated upon by medium (mdz) and large (lgz) zooplankton and higher trophic level predators with high preference (Fig. 1b).

While they have relatively large body sizes (e.g., the solitary form of *Salpa thompsoni* can exceed 10 cm; Dubischar et al., 2006), salps have fine mucous meshes that can filter submicron particles, and have a preference for grazing on small algae such as picoplankton (Sutherland et al., 2010). Salps are able to consume diatoms and bacteria, but with relatively low efficiency (Dadon-Pilosof et al., 2019) and with many diatoms passing through salp guts undigested (Harbison et al., 1986). Additionally, salps have been shown to be dominant grazers on *Trichodesmium* in certain regions (Post, 2002). While, the prey of doliolids is less studied, Walters et al. (2019) found using genetic approaches that doliolids preferentially consume diatoms (particularly in their early life stages) and ciliates. Thus, in the present model, the large tunicates (lgt) were able to feed on small phytoplankton (smp) and diazotrophs (diaz) with high preference (100% prey availability), microzooplankton (smz) at medium preference (50% prey availability), and bacteria (bact) and large phytoplankton (lgp) at low preference (25% prey availability).

Thaliacean predators and parasites were traditionally thought to be primarily sapphirinid copepods and large hyperiid amphipods, including the predatory and parasitic *Phronima* spp., which consume salp tissue and live in their cleared-out barrels (Laval, 1980; Madin and Harbison, 1977; Takahashi et al., 2013). Restricted emphasis on these relatively rare crustacean predators and parasites has yielded the misconception of gelatinous zooplankton, particularly salps, as trophic dead-ends. However, increasing evidence has highlighted the role of thaliaceans as food for fish and other higher trophic levels. Over 200 species of fish, turtles, corals, and echinoderms consume salps, doliolids, and pyrosomes, with many predators filling their guts with thaliaceans during bloom periods (Harbison, 1998; Henschke et al., 2016; Mianzan et al., 2001). Therefore, in our model, large tunicates (lgt) are predated upon by medium (mdz) and large (lgz) zooplankton at low preference (25% prey availability), reflecting the specialized nature of the copepod and amphipod predators relative to the broader crustacean zooplankton population, and by higher trophic level predators (hp) at medium preference (50% prey availability).

The diets of small and large tunicates are fairly similar in the model, as they are both microphagous generalists. Notably, there was no size scaling in the tunicates’ diets relative to their body size (i.e., smaller tunicates did not consume smaller prey), which is in contrast to crustacean mesozooplankton; large tunicates have larger predator-to-prey size ratios than small tunicates (Conley et al., 2018). Rather, the key distinction with respect to food web dynamics between the tunicates is the level of predation. Small tunicates (smt) have very strong levels of top-down control, exerted by all mesozooplankton (mdz and lgz) and higher trophic level predators (hp). While large tunicates (lgt) experience predation by similar predators as small tunicates in the model, the strength of that predation is reduced to account for their larger size. Other distinctions between the two tunicates, including ingestion rates, metabolic scalings, and susceptibility to jelly-falls are described in the next sections.

#### 2.2.2 Ingestion and assimilation

As filter feeders, the prey consumption rates of pelagic tunicates are typically measured as biomass-specific clearance (or filtration) rates, yielding ingestion rates (*I*, mg C m^-3^ d^-1^) as the product of the specific clearance rate (*C_b_*, m^3^ mg C^-1^ d^-1^), prey biomass (*P*, mg C m^-3^), and gelatinous predator biomass (*GZ*, mg C m^-3^)(Acuña et al., 2011):

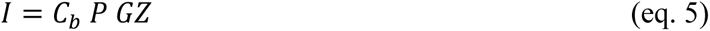

In contrast, COBALTv2 and other NPZD-type models utilize saturating functional response (Hollings Type II in the case of one prey) for zooplankton grazing (Fasham et al., 1990; Gentleman et al., 2003; Stock and Dunne, 2010), which is reproduced in eq. (3). When the ingestion half-saturation constant *K_i_* is much greater than the prey biomass (*K_i_* >> *P*), eq. (3) reduces to *I* = (*I_max_*/*K_i_*)\**P*\**GZ*, and *C_b_* becomes equivalent to *I_max_/K_i_* (Acuña and Kiefer, 2000). Therefore, we can use measured clearance rates to constrain the relationship between *I_max_* and *K_i_*.

For small tunicate clearance rates, we used allometric relationships from Lombard et al. (2009), which measured physiological rates for a common appendicularian, *Oikopleura dioica*. We assumed the small tunicates to be between 0.3-3 mm in body length, which yields a characteristic individual body length of 1 mm and associated biomass of 6.68 µg C. For a 6.68 µg C individual, specific filtration rates at 0°C (converted using a Q10 of 1.88) range from 0.010-0.017 (mean: 0.013) m^3^ mg C^-1^ d^-1^ (Lombard et al., 2009a). Considering that the clearance rates of both small and large tunicates do not significantly change with low to medium food concentrations (Gibson and Paffenhöfer, 2000; Paffenhöfer and Köster, 2011), we opted for a higher tunicate half-saturation constant (250 mg C m^-3^) than that of crustacean zooplankton (102 mg C m^-3^; Table 1), consistent with the estimated range of *K_i_* for *O. dioica* of 20-500 mg C m^-3^ (Acuña and Kiefer, 2000). The equivalent *I_max_* would be 2.50-4.25 (mean: 3.25) d^-1^. Following model tuning, we used an *I_max_* value on the low end of the range (1.875 d^-1^) given a mean *K_i_*, to avoid overconsumption by small tunicates. However, considering the wide variation in *K_i_*, these values were well within the observational bounds. The tradeoffs between tunicates and crustacean zooplankton were visualized in a plot of specific ingestion at 25°C (Fig. 2a): the small tunicate *I_max_* and *K_i_* results in a specific ingestion in between small/micro-zooplankton and medium/crustacean zooplankton. Additionally, a sensitivity run was conducted to illustrate the effect of the *I_max_* tuning choice (Section 2.3).

**Figure 2.**
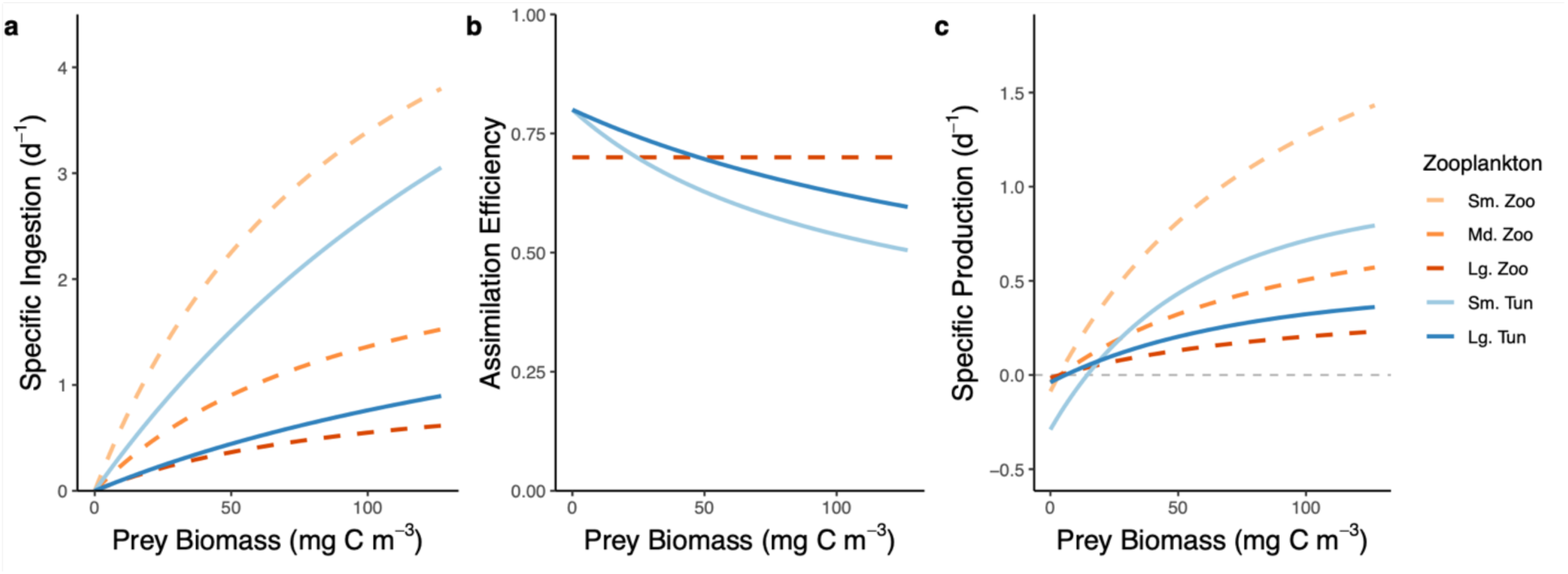
Specific ingestion (a), assimilation efficiency (b), and specific production (assimilated ingestion minus respiration) (c) as a function of generalized prey biomass at 25°C for zooplankton in GZ-COBALT. The three original COBALTv2 zooplankton types (small, medium, and large zooplankton; dashed lines) are unchanged in all GZ-COBALT simulations from the COBALTv2 control model.

For large tunicates, we were able to use an allometric scaling relationship from a prior effort (Luo et al., 2020) that compiled length, carbon biomass, and clearance rate relationships from various studies (see Madin and Deibel, 1998) into a single equation. We assumed the large tunicates spanned 3-300 mm in body length, which yields a characteristic individual body length of 28 mm and associated biomass of 1.5 mg C. At this biomass, specific clearance rates at 0°C range from 4.2E-4 to 7.4E-3 (mean: 1.8E-3) m^3^ mg C^-1^ d^-1^. Using a *K_i_* of 250 mg C m^-3^, and the same relationship between clearance rates, *K_i_*, and *I_max_* as above, we estimated large tunicate *I_max_* values to be between 0.105-1.85 d^-1^. In the model, we used a value in the lower half of the reported range (0.55 d^-1^), tuned in conjunction with other variables with wide uncertainty bounds to match observed biomass concentrations. At the lowest prey concentrations, the *I_max_/K_i_* of large tunicates matched that of the large crustacean zooplankton. As prey concentrations increased, large tunicate ingestion fell between that of the medium and large crustacean zooplankton and reached its maximum ingestion rate much slower than either crustacean group (Fig. 2a).

Assimilation efficiency (*AE*) for zooplankton is typically set to be a fixed fraction of ingested material in models (between 0.6-0.8; Carlotti et al., 2000), and for crustacean zooplankton in COBALTv2, it is set to 0.7, allowing for 70% of all food consumed to be assimilated independent of prey concentration. However, for pelagic tunicates, in particular appendicularians, there is evidence of *AE* declines as prey concentration increases, due to development of tears of their pharyngeal filter and active prey rejection with increasing food (Acuña and Kiefer, 2000; Lombard et al., 2009a, 2011). Retention and assimilation efficiencies for salps and doliolids also vary widely, from 28-90%, which may be due to prey selectivity for optimal sizes and preferred taxa (Andersen, 1986; Dadon-Pilosof et al., 2019; Pakhomov, 2004; Pakhomov et al., 2006; Vargas and Madin, 2004; Walters et al., 2019). While we have implemented prey selectivity in the feeding relationships, there is still evidence for feeding apparatus clogging at high food concentrations due to the formation of a food bolus (Harbison et al., 1986). Therefore, we implemented varying assimilation efficiencies for pelagic tunicates:

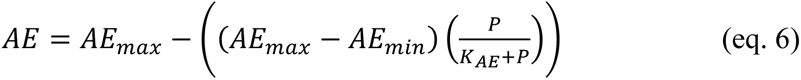

where *AE_max_* and *AE_min_* are maximum and minimum assimilation efficiencies, respectively (unitless), and *K_AE_* is the half-saturation constant for AE (mg C m^-3^).

This AE equation is a desaturating functional form (Hollings Type II subtracted from *AE_max_*)(see also Berline et al., 2011). For small tunicates, *AE_max_* and *AE_min_* were 0.7 and 0.25, respectively, and *K_AE_* was 110 mg C m^-3^, which is at the low end of the range for the appendicularian *O. dioica* (145.52 +/- 33.36 std. err.) as measured by Lombard et al. (2009a). For large tunicates, to simulate the clogging response, we also used the same *AE* bounds as small tunicates but with a *K_AE_* of 215 mg C m^-3^, a value at which approximately half of the *Pegea confoederata* salps studied by Harbison et al. (1986) would form boluses. The difference in *K_AE_* between the tunicates results in the *AE* declining faster for small tunicates compared to their larger counterparts (Fig. 2b).

Non-assimilated egestion losses for tunicates were parameterized similar to large zooplankton with 100% of the detritus losses going towards sinking particulate organic matter and no dissolved organic matter generated through grazing. This assumption is consistent with the representation of small tunicates in Berline et al. (2011) and the observation that tunicate detritus, representing the sinking houses of appendicularians and salp fecal pellets, are an important source of zooplankton detritus (Alldredge and Silver, 1988), and contribute significantly to carbon export fluxes (Alldredge, 2005; Andersen, 1998; Luo et al., 2020; Robison, 2005).

In GZ-COBALT, we did not modify the sinking speed of GZ-mediated detritus, which is pooled together with non-gelatinous detritus and sinks at a rate of 100 m d^-1^ (Alldredge and Silver, 1988; Stock et al., 2014a; Turner, 2015). While studies have shown that GZ detritus can sink at rates greatly exceeding that of marine aggregates and crustacean zooplankton fecal pellets (Lebrato et al., 2013), we opted to focus this study on the impact of gelatinous zooplankton, specifically tunicates, within the euphotic zone and leave the assessment of GZ-mediated export on biogeochemical cycles at depth to future work.

#### 2.2.3 Metabolism and respiration

In the COBALTv2 base model, the total zooplankton respiration rate is a sum of the basal (*bresp*) and active (*aresp*) respiration rate, with the former the resting metabolic rate that is proportional to biomass and the latter a fixed fraction of the ingestion rate (Flynn, 2005; Stock et al., 2014a). Pelagic tunicates are true filter feeders, and in contrast with crustacean zooplankton, are constantly in motion regardless of the food concentration, pumping water via tail oscillations (appendicularians), muscle contractions (salps), and ciliary action (doliolids and pyrosomes) (Deibel, 1998; Madin and Deibel, 1998). Therefore, a truly “basal” respiration rate only occurs when the tunicate is anesthetized. For *Salpa fusiformis*, swimming accounts for over half of total oxygen demand (Trueman et al., 1984), while for the appendicularian *O. dioica,* swimming accounted for roughly 34% of active respiration (Lombard et al., 2005). Furthermore, Lombard et al. (2005) found that there was no significant difference between the respiration levels of appendicularians at various food levels (no food, low, and high food), implying that the energy required to digest food was small compared to the energetic requirements of swimming. In contrast, the basal respiration rates for crustacean zooplankton are relatively low compared to respiration rates while feeding, which increase linearly with ingestion (Kiørboe et al., 1985). Thus, a key contrast that is built into GZ-COBALT is the difference between crustacean and gelatinous zooplankton respiration tradeoffs: crustaceans have relatively high active respiration rate (30% of ingestion) but low basal respiration, whereas GZ have low active respiration (15% of ingestion) but high basal respiration rates (Table 1). Consequently, compared to crustacean zooplankton, tunicates incur a higher metabolic cost in low food concentrations, which prevents them from accumulating biomass in large portions of the subtropical gyres. These tradeoffs can be seen on a plot of specific production as a function of food concentration (Fig. 2c), and is most obvious when comparing small tunicates vs. small mesozooplankton, as the tunicate specific production remains negative for a greater portion of the low food concentrations. A full comparison of parameters is found in Table 1.

Biomass-specific basal respiration for small tunicates (smt) was calculated using relationships for *O. dioica* (Lombard et al., 2005). A laboratory study in the absence of food found that a 6.68 μg C individual at 0°C has a weight-specific oxygen consumption of 0.068 (0.055-0.085) µmol O_2_ (µmol C)^-1^ d^-1^ (Lombard et al. 2005). Assuming a general zooplankton respiratory quotient of 0.87 (Mayzaud et al., 2005), this translates to a basal respiration of 0.047-0.074 d^-1^. We opted to use a value at the low end of the range (0.047 d^-1^), so as to avoid complete elimination of their biomass in subtropical environments, where they are found in low concentrations (Steinberg et al., 2008).

For large tunicates (lgt), GZ-COBALT takes advantage of the mean allometric respiration relationship compiled by Luo et al. (2020) from observations in Madin and Deibel (1998; and references therein). With an average large tunicate of 1.5 mg C in that study respiring 2.5E-3 (8.0E-4 – 8.1E-3) mmol O_2_ mg C^-1^ d^-1^, and using a salp-specific respiratory quotient of 1.16 (Mayzaud et al., 2005), basal respiration varied by an order of magnitude (0.011-0.11 d^-1^) with mean of 0.035 d^-1^. However, even with this large uncertainty range, Luo et al. (2020) found that under average conditions in the pelagic oceans, even the lower bound of these respiration rates were too high, such that metabolic demands exceeded available food resources, yielding unfavorable conditions for survival, particularly in the subtropical gyres. Therefore, in the baseline model configuration, we used a basal respiration rate (0.008 d^-1^) that was slightly below the lower bound, tuned to ensure realistic gelatinous zooplankton in oligotrophic ecosystems. This choice is consistent with the strategy enlisted for calibrating the basal metabolic costs of the crustacean zooplankton (Stock and Dunne, 2010) wherein highly uncertain basal metabolic rates were calibrated to ensure that the simulated biomass was consistent with observations in the oligotrophic gyres where the impact of basal metabolic costs are most prominent. To explore the effect of this tuning, a sensitivity case was run where large tunicate basal respiration was set to the mean value from allometry (Section 2.3).

#### 2.2.4 Other sources of mortality

Another key difference between the small and large tunicates is in the additional sources of mortality. While small tunicates primarily experience mortality through predation, large tunicates experience cold temperature reproductive failures (Henschke and Pakhomov, 2019) as well as mass mortality events (jelly-falls) (Henschke et al., 2015; Lebrato and Jones, 2009). Large tunicate (lgt) aggregation losses, representing jelly-falls, were parameterized as a quadratic loss that is suppressed when food is plentiful, following the same functional form as the phytoplankton aggregation losses as a function of nutrient limitation in COBALTv2 (Stock et al., 2020; Waite et al., 1992) and other global biogeochemical models (e.g., PISCES; Aumont et al., 2015). The aggregation loss (*agg_lgt_*) is controlled by two parameters: *f_agg_,* which represents the fraction of the maximum ingestion rate above which aggregation losses are suppressed, and *m_agg_,* or an aggregation loss rate. The former parameter is used to set a growth ratio (*µ_ratio_*) that calculates the total ingestion relative to a fraction of the maximum ingestion:

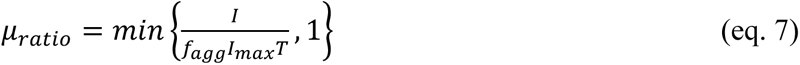

Then, aggregation loss is a density-dependent term that decreases to zero when µ_ratio_ = 1 and increases quadratically as µ_ratio_ < 1:

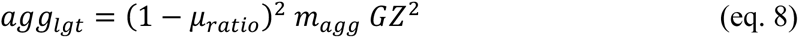

We set *f_agg_* to 0.l to account for the salp ability to tolerate low food concentrations, and *m_agg_* to 1.0E-3 m^3^ mg C^-1^ d^-1^ to achieve jelly-falls representing approximately 35% of total large tunicate mortality, following Luo et al. (2020).

### 2.4 Physical Framework

The GZ-COBALT model with 35 tracers was run in a global ocean-ice configuration using the GFDL models Modular Ocean Model 6 (MOM6) and Sea Ice Simulator 2 (SIS3) in a nominal 0.5°, or roughly 50 km, horizontal resolution (OM4p5, Adcroft et al., 2019). The 0.5° horizontal grid improves the resolution of boundary currents compared to earlier generations of 1° MOM models. The vertical coordinate in MOM6 is a hybrid z*-isopycnal vertical coordinate system implemented using an Arbitrary Lagrangian-Eulerian (ALE) method, such that isopycnal coordinates are used in the ocean interior and a z* coordinate is used in the mixed layer. OM4p5 uses 75 vertical layers, which allows for finer resolution at the ocean surface (∼2 m) compared to earlier model configurations with 10 m surface resolution and 50 vertical layers (Adcroft et al., 2019). The ocean and ice model configurations are also equivalent to those components used within the fully-coupled ESM4.1 model (Dunne et al., 2020).

Model simulations were forced using Common Ocean-Ice Reference Experiment II (CORE-II) (Large and Yeager, 2009), a 60-year interannually varying dataset representing atmospheric forcings from 1948-2007. The model was initialized similar to that of the fully-coupled model (Stock et al., 2020): from World Ocean Atlas 2013 (WOA13) data for temperature, salinity, oxygen, and dissolved inorganic nutrients (Garcia et al., 2013a, 2013b; Locarnini et al., 2013; Zweng et al., 2013), and from Global Ocean Data Analysis Project (GLODAPv2) for dissolved inorganic carbon and alkalinity (Lauvset et al., 2016). Other tracers were initialized from outputs of a previous version of COBALT (Stock et al., 2014a), and the two new gelatinous zooplankton tracers were initialized with biomass concentrations similar to medium and large zooplankton. Additional sources of nutrients include atmospheric deposition of NH_4_ and NO_3_ (Horowitz et al., 2003), dust from Zhao et al. (2018) with soluble Fe calculated in accordance with Baker and Croot (2010), as well as coastal iron and river nutrients from the GlobalNEWS dataset (Seitzinger et al., 2005), as described in (Stock et al., 2020)

GZ-COBALT was run for one 60-year interannual forcing cycle. Results are reported from a climatology computed from the last 20 years of model simulation, representing 1988-2007. A COBALTv2 control simulation with the same exact model setup, but with tunicates turned off, was also run for 60 years.

### 2.5 Parameter Sensitivity Runs

To understand the impact of the unique aspects of GZ physiology and ecology as described above on their emergent distribution and productivity, we considered a number of perturbations around the baseline settings described above. These sensitivity runs (Table 2) examine both the effect of our tuning choices (cases 1-2) as well as some key tradeoffs relative to crustacean zooplankton (cases 3-5). All parameter sensitivity runs were conducted following the same physical forcing as the GZ-COBALT base simulation.

1) For large tunicates, the basal respiration rate was adjusted to be slightly below the lower bound from the literature. Sensitivity case 1 examines the effects of using the mean basal respiration rate from allometric relationships from the literature.
2) For the baseline GZ-COBALT model, we used a maximum ingestion rate of small tunicates at the lower bound of the range. For sensitivity case 2, we test a case where the small tunicates’ I_max_ is higher, set to the mean of the literature-based range.
3) Sensitivity case 3 explores the impact of the unique feeding behavior of GZ relative to crustaceans. For the calibrated model (base case), we used a mean K_i_ value from a wide range measured by Acuña and Kiefer (2000). However, models are known to be quite sensitive to this relatively unconstrained parameter (Stock and Dunne, 2010). Thus, we ran a sensitivity test where the tunicate K_i_ values were the same as that of the crustacean zooplankton. Since the maximum ingestion rate (I_max_) was set in combination with the K_i_ values to achieve specific filtration rates at low prey concentrations consistent with observations, I_max_ was also modified accordingly to preserve the observational constraint.
4) Sensitivity case 4 examines another unique aspect of pelagic tunicates feeding relative to that of crustaceans. In the baseline GZ-COBALT, we implemented varying assimilation efficiency (AE) following Berline et al. (2011). To explore the uncertainty associated with this assumption, we ran a sensitivity test where the tunicates’ AE were constant, but set at a slightly lower value than other zooplankton, to account for their comparatively lower retention rates.
5) Finally, a case was run to explore the impacts of ignoring the role of large tunicate aggregation mortality (representing jelly-falls).

**Table 2.**
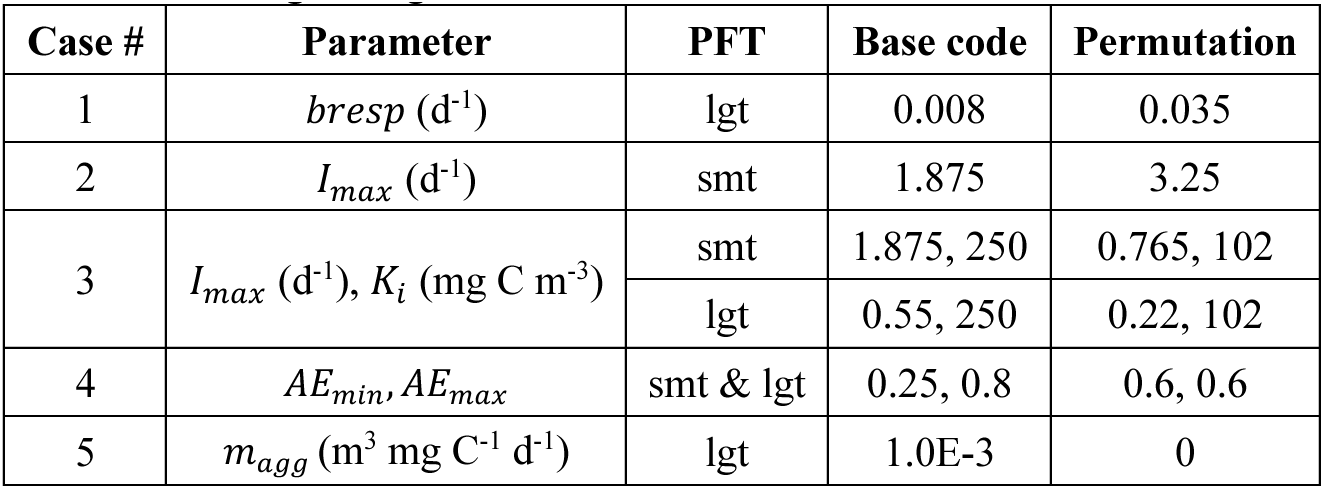
Parameters tested in sensitivity tests, showing the values in the base code as well as the permutation for the individual sensitivity case. For detailed names of parameters, see Table 1. Plankton functional types (PFTs): smt = small tunicates; lgt = large tunicates.

| Case # | Parameter | PFT | Base code | Permutation |
| --- | --- | --- | --- | --- |
| 1 | $bresp$ ($d^{-1}$) | lgt | 0.008 | 0.035 |
| 2 | $I_{max}$ ($d^{-1}$) | smt | 1.875 | 3.25 |
| 3 | $I_{max}$ ($d^{-1}$), $K_i$ ($mg\ C\ m^{-3}$) | smt | 1.875, 250 | 0.765, 102 |
|  |  | lgt | 0.55, 250 | 0.22, 102 |
| 4 | $AE_{min}$, $AE_{max}$ | smt & lgt | 0.25, 0.8 | 0.6, 0.6 |
| 5 | $m_{agg}$ ($m^3\ mg\ C^{-1}\ d^{-1}$) | lgt | 1.0E-3 | 0 |

To illustrate the impact of these various parameter modifications, we computed the specific production (assimilated ingestion minus respiration) of all zooplankton groups under an idealized condition of 25°C with a generalized prey biomass (Fig 3), which can be contrasted against the GZ-COBALT baseline (Fig. 2c).

**Figure 3.**
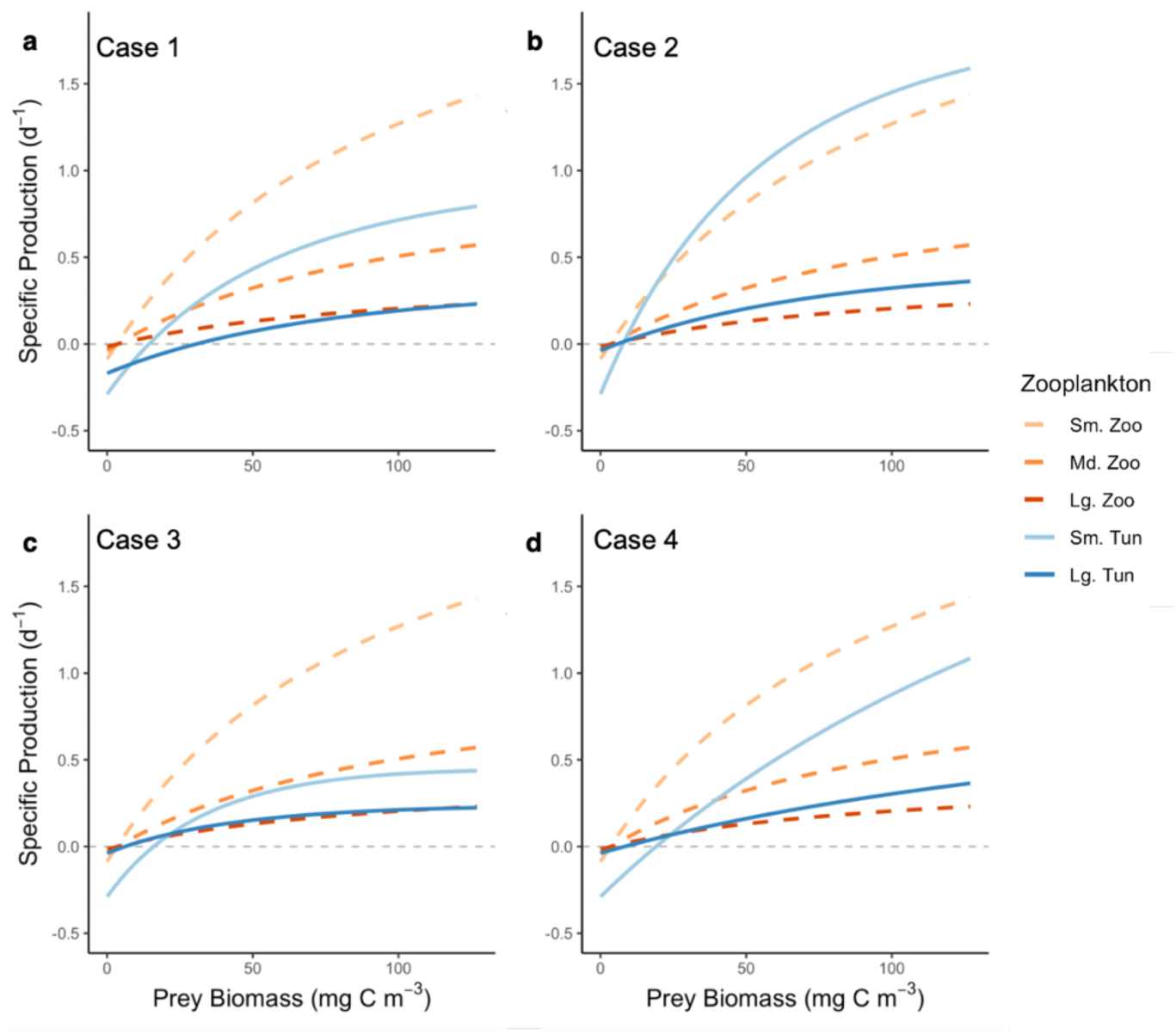
Specific production as a function of generalized prey biomass at 25°C for sensitivity cases 1-4 (a-d). The three original COBALTv2 zooplankton types (small, medium, and large zooplankton; dashed lines) are unchanged in all GZ-COBALT simulations from the COBALTv2 control. The specific production in sensitivity case 5 is the same as the base case (Fig. 2c).

### 2.6 Validation data

As described in the introduction, a central question in our analysis is whether the model can simultaneously reconcile recent measurements suggesting that pelagic tunicate carbon biomass is *ca.* 10x greater than previously thought (cf. Remsen et al., 2004) while also satisfying core observational constraints on crustacean biomass, nutrients, chlorophyll and net primary production (NPP). The subsections that follow describe the datasets compiled and/or enlisted for each of these tasks, and the analyses used to assess model-data consistency across trophic gradients and ocean biomes.

#### 2.6.1 Pelagic tunicate dataset and model-data comparison

To generate biomass validation data for pelagic tunicates, we updated the gridded tunicate data (primarily salps) from Luo et al. (2020) with additional data on both thaliaceans and appendicularians from the NOAA Coastal and Oceanic Plankton Ecology, Production, and Observation Database (COPEPOD) (O’Brien, 2014). From the COPEPOD database, raw data on urochordates were extracted and divided into small tunicates (all appendicularians) and large tunicates (salps, doliolids, and pyrosomes). With the exception of data from the ECOSAR-II cruise (Muxagata, 1999), all other data were in numeric density only (# individuals m^-3^). Numeric density data were first converted to a common 330 µm mesh size (Moriarty and O’Brien, 2013; O’Brien, 2005). Second, since the geometric mean cannot handle zeros, zero numeric density values were modified to be a non-zero value slightly below the minimum value for both size fractions (small: 0.0008 ind. m^-3^, large: 0.001 ind. m^-3^). Next, numeric density was converted to carbon biomass using the characteristic individual biomass values defined in section 2.2.2 (appendicularians: 6.7 µg C, salps: 1.5 mg C). Characteristic biomass values for pyrosomes and doliolids were 22.9 mg C ind^-1^ (100 mm individual) and 19.2 µg C ind^-1^ (5 mm individual), respectively, following Lucas et al. (2014), which used regression conversions from Gibson and Paffenhöfer (2000) and Mayzaud et al. (2007). Using the geometric mean, 1° gridded values were averaged by month, then year, for an annual mean biomass. Finally, the Jellyfish Database Initiative (JeDI; Condon et al., 2015) database was additionally queried for appendicularian data. Data from 90 additional 1° grid cells, primarily from the North Atlantic and Eastern Equatorial Pacific, were present in the JeDI dataset but not in the COPEPOD database. These data were added to our validation dataset. Appendicularian data were present in a total of 3,914 1° grid cells (Fig. 4a).

**Figure 4.**
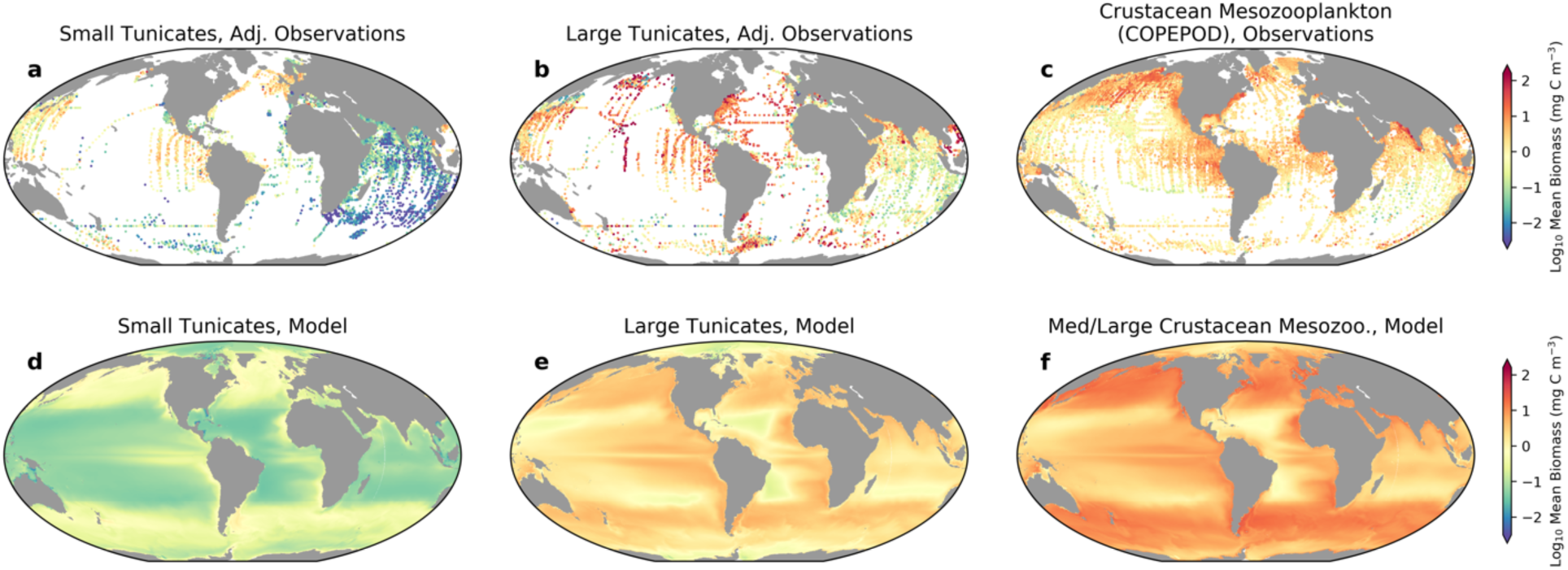
Global mean distributions of small and large tunicates and crustacean mesozooplankton, comparing tunicate adjusted observations (a,b), large crustacean biomass from COPEPOD (c) with results from the top 100 m of the model (d,e,f). Medium/large crustacean mesozooplankton model values are given as the large mesozooplankton plus 0.5*small mesozooplankton, to reflect the fact that the net sizes in the COPEPOD database largely do not capture mesozooplankton in the smaller size ranges. Model data show the time-average of the growing season only (fall and winter months excluded poleward of 30°N/S).

Thaliacean data from the COPEPOD database were combined with the Luo et al. (2020) gridded salp data, which primarily included gridded biomass data from Lucas et al. (2014), with updates from JeDI, the Palmer LTER site at the Western Antarctic Peninsula (Steinberg et al., 2015), and KRILLBASE (Atkinson et al., 2017). Out of the 5,468 grid cells with data, there were 1,481 cells where COPEPOD data were only present, 1,952 cells where the Luo et al. (2020) data were only present, and overlap at 2,035 grid cells (Fig. 4b). Because the raw data with assumed lengths and carbon conversions from Lucas et al. (2014) were not available, we were unable to examine individual data points for overlap and cross-validation. However, a broad examination of the two datasets revealed that at the areas of overlap, carbon biomass compiled from COPEPOD was 1.4x (geometric mean) that of Luo et al. (2020), with variations likely due to the finer taxonomic detail of the Lucas et al. (2014) effort. This was within the uncertainty bounds that we considered acceptable (roughly 2x uncertainty). Ultimately, due to discrepancies in classification and specificity over time (e.g., broad categories such as “Tunicata” and “Salps and doliolids” were dominant in classifications from the 1950’s and 1960’s, but not later), we decided that using a single characteristic carbon biomass conversion for each broad taxonomic category in the COPEPOD data gave a taxonomic specificity consistent with the coarsest taxonomic specificity in the data. Further, this single biomass conversion removed a persistent discontinuity in the Luo et al. (2020) carbon biomass values in the Indian Ocean south of 5°N that we were previously unable to resolve. Since the vast majority of the JeDI data sources in the Indian Ocean were from the 1959-1965 International Indian Ocean Expedition (IIOE), which are also present in COPEPOD (Condon et al., 2015), we opted to replace the Luo et al. (2020) Indian Ocean data with the COPEPOD data during our merge. For the rest of the oceans, we merged the two datasets by taking the geometric mean at every grid cell where there was overlap.

Ultimately, the discrepancies between the datasets were quite low compared to the differences in biomass due to sampling type, particularly when comparing extractive (nets) vs. non-extractive (imaging) methods. Since traditional, net-based sampling systems break apart fragile organisms such as pelagic tunicates and other gelatinous zooplankton (and comprise the vast majority of the data in this present compilation), a biomass adjustment is necessary to account for the reduced sampling from nets. Remsen et al. (2004) used concurrent sampling with an imaging system and a 162 µm mesh plankton net, and found that for pelagic tunicates, their abundance was undersampled by nets by a factor of 3-4x, and their carbon biomass undersampled by a factor of 5-15x. Therefore, we considered an additional “adjusted biomass” from samples with a 10x increase relative to the unadjusted biomass. We focus mainly on this adjusted biomass, which is indicative of nascent appreciation of the likely broader importance of gelatinous zooplankton as revealed by optical instruments. This is also consistent with our intent to assess whether these high values can be reconciled with the overall high abundance of mesozooplankton in many regions.

We complement our assessment of the simulated magnitude of tunicate biomass with one of the relationship between tunicate biomass and other ecosystem properties spanning oceanographic gradients. As the spatial gradients in tunicate biomass span 5 orders of magnitude, this assessment provides a second metric less sensitive to the adjustments above. To do this, we considered the GZ biomass as a function of chlorophyll concentration. The resultant, large-scale relationship allowed for contrasts between large and small tunicates, and between tunicates and crustaceans. For chlorophyll, we used the GlobColour merged satellite chlorophyll product (from MERIS, MODIS-Aqua, and SeaWiFS) monthly climatology for case 1 waters using the weighted averaging method, blended at latitudes south of 50°S with the Southern Ocean algorithm of Johnson et al. (2013). We computed a growing season mean, define as all months for latitudes between 30°N and 30°S, and spring and summer only for latitudes poleward of 30°N/S. The slope of the log-log relationships between chlorophyll and the biomass of small tunicates, large tunicates, and crustacean mesozooplankton (Moriarty and O’Brien, 2013; more details in Section 2.6.3) were established as emergent relationships for validation purposes.

#### 2.6.2 Biome definitions

Finally, we assess gelatinous zooplankton simulation as a function of ocean biome, adjusting for any systematic biases in the model by referencing biome locations to chlorophyll, light, and temperature thresholds. We used the three major ocean biomes of Stock et al. (2014), following Banse (1992): 1) Low Chlorophyll (LC), which encompasses the subtropical gyres, 2) High Chlorophyll Seasonally Stratified (HCSS), which encompasses the high latitudes, and 3) High Chlorophyll Permanently Stratified (HCPS), which includes the coastal and equatorial upwelling regions. Stock et al. (2014) used a threshold of 0.125 mg Chl m^-3^ to separate between the low vs. high chlorophyll regions in observational chlorophyll datasets. In our biome definition, we first calculated the total ocean area with observational chlorophyll values lower than that threshold (approximately 40% of the world’s oceans), then found the model chlorophyll threshold that resulted in a model LC area that most closely matched the LC surface area from observations. For the COBALTv2 control and GZ-COBALT, this threshold was 0.162 and 0.184 mg Chl m^-3^, respectively. Next, to distinguish between the seasonally vs. permanently stratified regions, we used the minimum of the mixed layer irradiance climatology (light averaged over the mixed layer). HCSS regions were demarcated as those with minimum mixed layer irradiances lower than 5 W m^-2^, while the opposite was true of HCPS. Using mixed layer irradiance more accurately defined the seasonal seas vs. upwelling areas, preventing HCPS areas from occurring in Arctic regions with shallow maximum mixed layers (Stock et al., 2014a). Biomes for both GZ-COBALT and the COBALTv2 control are shown in Fig. S1.

#### 2.6.3: Crustacean zooplankton dataset and model-data comparison

To assess whether pelagic tunicate biomass magnitude and cross-biome gradients can be represented while maintaining crustacean zooplankton populations consistent with observations, we used the 2012 gridded carbon biomass data compilation from the COPEPOD database (Moriarty and O’Brien, 2013). The entire COPEPOD database (O’Brien, 2014) consists of multiple types of data products, including the raw, taxonomic data as used above for pelagic tunicates, as well as the Moriarty and O’Brien (2013) carbon biomass compilation, which is the more commonly used dataset for mesozooplankton model validation. In total, the COPEPOD global carbon biomass compilation includes over 150,000 data points that were converted to an equivalent 333 µm mesh net size, with each gridded value representing multiple data points. Given the net-based sampling, the mesh size, and the historical focus on crustacean zooplankton, the vast majority of the individual data points consisted of relatively large, hard-bodied mesozooplankton. Thus, we used the COPEPOD global carbon biomass compilation as a proxy of the medium to large crustacean mesozooplankton, which can be compared against the large crustacean mesozooplankton (lgz) plus half of the small crustacean mesozooplankton (0.5*mdz) in GZ-COBALT. We did not use the full crustacean mesozooplankton biomass field as the COPEPOD database greatly underestimates mesozooplankton in the smaller (200-500 µm) size ranges.

Similar to the GZ data, the crustacean observations were also scaled with chlorophyll on a log-log scale. This enabled us to make comparisons along trophic gradients and across biomes for crustaceans and gelatinous zooplankton.

Finally, in addition to GZ and crustacean constraints, we include a suite of standard biogeochemical metrics to ensure that the model solution satisfies large-scale productivity and nutrient patterns. The data we used were the dissolved inorganic nutrient concentrations (NO_3_, PO_4_, and SiO_3_) at the ocean surface from the World Ocean Atlas (WOA) 2018 (Garcia et al., 2019).

## 3. Results

### 3.1 Global distribution and biomass-chlorophyll scaling

The GZ-COBALT simulation produced mean values consistent with the adjusted biomass of small and large pelagic tunicates, while also reproducing observed crustacean biomass and satisfying ocean biogeochemical constraints (Figs. 4-5, Tables 3-4). Global NPP was 53.7 Pg C y^-1^ and export flux at 100 m was 6.36 Pg C y^-1^ in GZ-COBALT, compared to 55.4 and 6.23 Pg C y^-1^ in COBALTv2. Surface chlorophyll and macronutrient concentrations in GZ-COBALT also compared well with the COBALTv2 control and observational constraints (Fig. 5).

**Figure 5.**
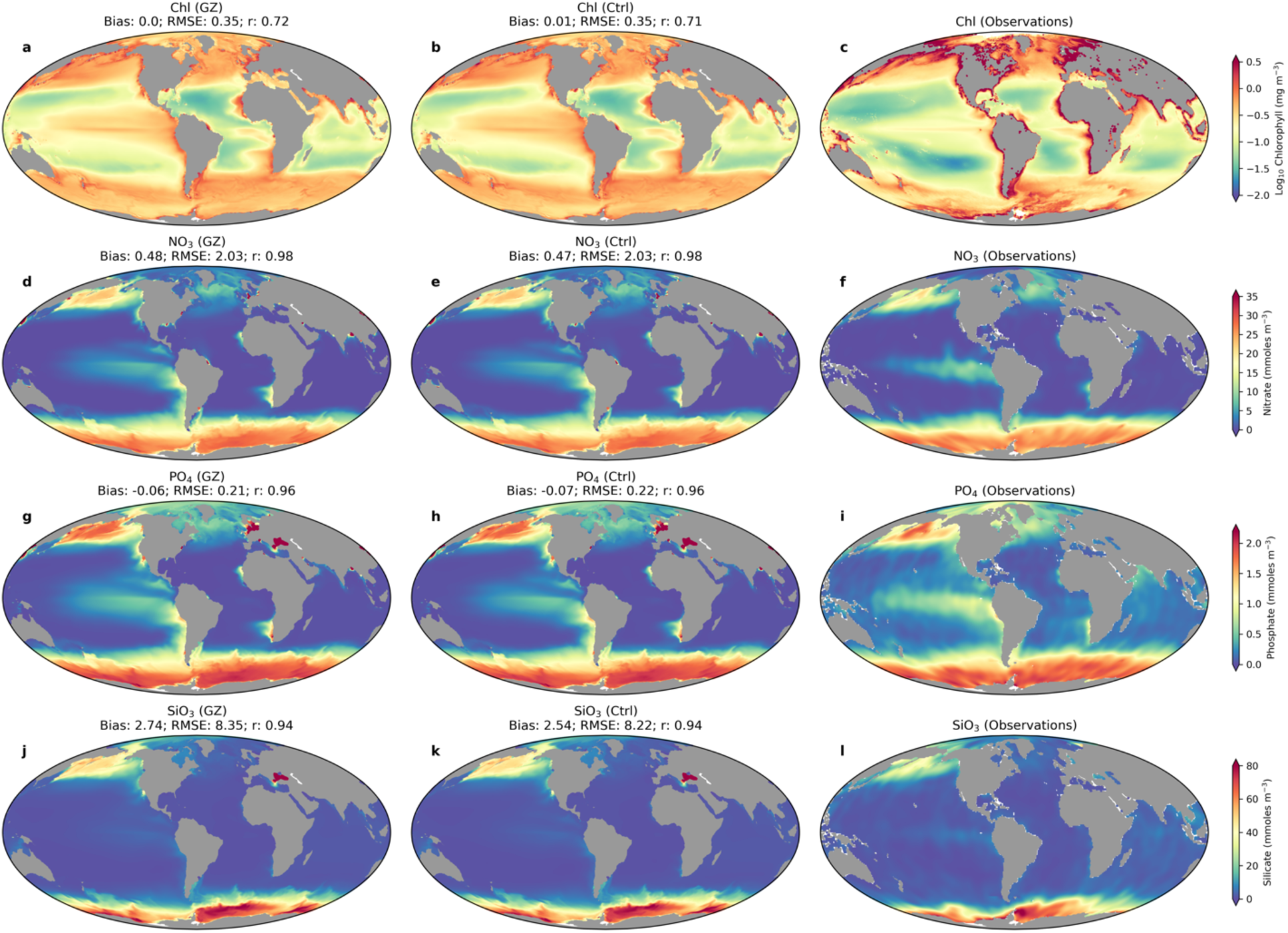
Surface chlorophyll, log10 scale (mg Chl m^-3^, a-c) and nutrients (mmol NO3 m^-3^, d-f; mmol PO4 m^-3^, g-i; mmol SiO3 m^-3^, j-i) from GZ-COBALT (left column; a, d, g, j), the COBALTv2 Control (center column; b, e, h, k), and observations (right column; c, f, i, l). Model bias, root mean squared error (RMSE), and Pearson’s correlation coefficient (r) are also reported.

**Table 3.** Global and biome-specific biomass comparison of the observations and the GZ-COBALT model. Model results are the annual area weighted mean of the top 100 m carbon biomass. For the medium to large crustacean mesozooplankton we used model fields lgz + mdz*0.5 to compare against the COPEPOD database. Observational values are given as the geometric mean and geometric standard deviation of values within the biomes, with global geometric standard deviation calculated following the procedure in Luo et al. (2020) and detailed in the SI text. Biomes: LC = Low Chlorophyll; HCPS = High Chlorophyll Permanently Stratified; HCSS = High Chlorophyll Seasonally Stratified. See Fig. S1, S2 for biome maps.

|  | Large Tunicates (mg C m<sup>-3</sup>) |  | Small Tunicates (mg C m<sup>-3</sup>) |  | Med/Large Crustacean mesozooplankton (mg C m<sup>-3</sup>) |  |
| --- | --- | --- | --- | --- | --- | --- |
| Biome | Adj. Obs. mean (stdev) | Model annual mean | Adj. Obs. mean (stdev) | Model annual mean | Obs. mean (stdev) | Model annual mean |
| LC | 1.4 (10) | 1.4 | 0.05 (16) | 0.06 | 0.98 (2.3) | 2.6 |
| HCPS | 1.9 (8.0) | 3.2 | 0.11 (15) | 0.16 | 3.6 (2.5) | 6.6 |
| HCSS | 2.9 (11) | 2.4 | 0.08 (23) | 0.28 | 2.9 (3.2) | 8.3 |
| **Global** | 2.0 (9.9) | 1.6 | 0.07 (18) | 0.11 | 2.0 (3.1) | 4.1 |

**Table 4.**
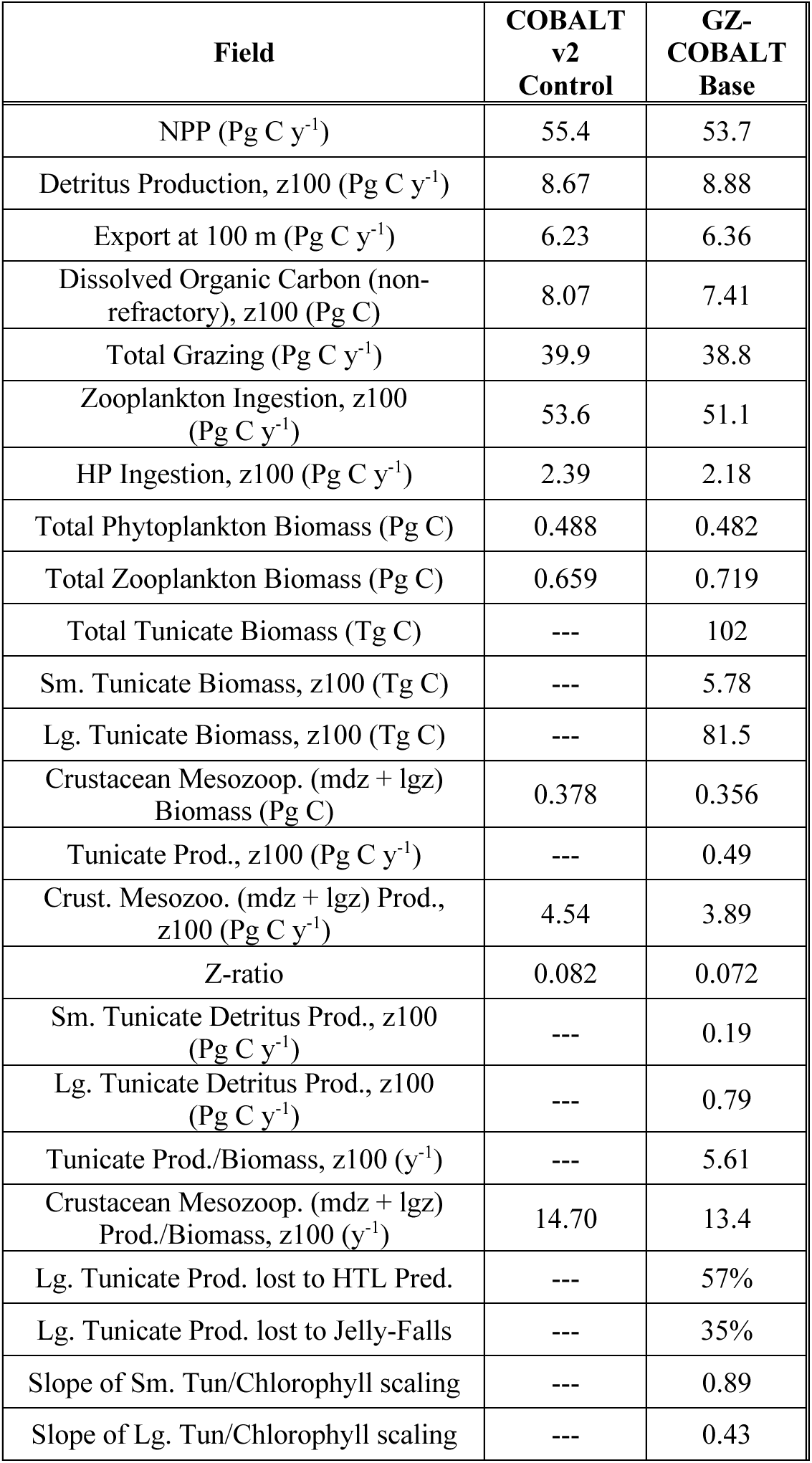
Comparison of the major results from the COBALTv2 and the GZ-COBALT base simulation. ‘z100’ refers to the top 100 m of the water column.

The modeled global mean annual biomass integrated over the top 100 m was 5.8 Tg C for small tunicates and 81.5 Tg C for large tunicates, yielding a total 100 m biomass of 87 Tg C. A small but non-negligible fraction of tunicate biomass was below 100 m, even with the model lacking vertical migration, such that the water column integrated biomass was 102 Tg C. These values are within the adjusted mean and uncertainty of the observations. In comparison, the medium/large crustacean mesozooplankton biomass (representing the size fraction most closely comparable to the values in COPEPOD database) in GZ-COBALT was 205 Tg C in the top 100 m, which was slightly lower than the COBALTv2 value of 220 Tg C. Observational estimates of large mesozooplankton biomass from COPEPOD, using a biome-specific geometric mean and standard deviation to extrapolate globally, was 133 (+/- 209) Tg C over the top 200 m. See Table 3 for additional comparisons by major ocean biome.

In the observations, we found that there was a contrast in the slope and intercept of biomass-chlorophyll scaling relationship between small tunicates, large tunicates, and crustacean mesozooplankton (Fig. 6). The small tunicates had significantly less biomass and a steeper log-log slope (0.63 +/- 0.045 residual std. err.; Fig. 6a) than the large tunicates, which was much flatter (0.22 +/- 0.036; Fig. 6b). The crustacean mesozooplankton data had much less variability, a mean biomass similar to that of the large tunicates, and a biomass-chlorophyll scaling slope a little shallower than the small tunicates (slope: 0.57 +/- 0.009; Fig. 6c). GZ-COBALT successfully captured the differences in mean biomass across all three groups, as well as the contrast in slope between the three groups, though admittedly the modeled slopes were all slightly steeper than the observational slopes. The large tunicates had the shallowest biomass-chlorophyll scaling slope (0.43, Fig. 6b), followed by the crustacean mesozooplankton (0.71, Fig. 6c) and the small tunicates (0.89, Fig. 6a).

**Figure 6.**
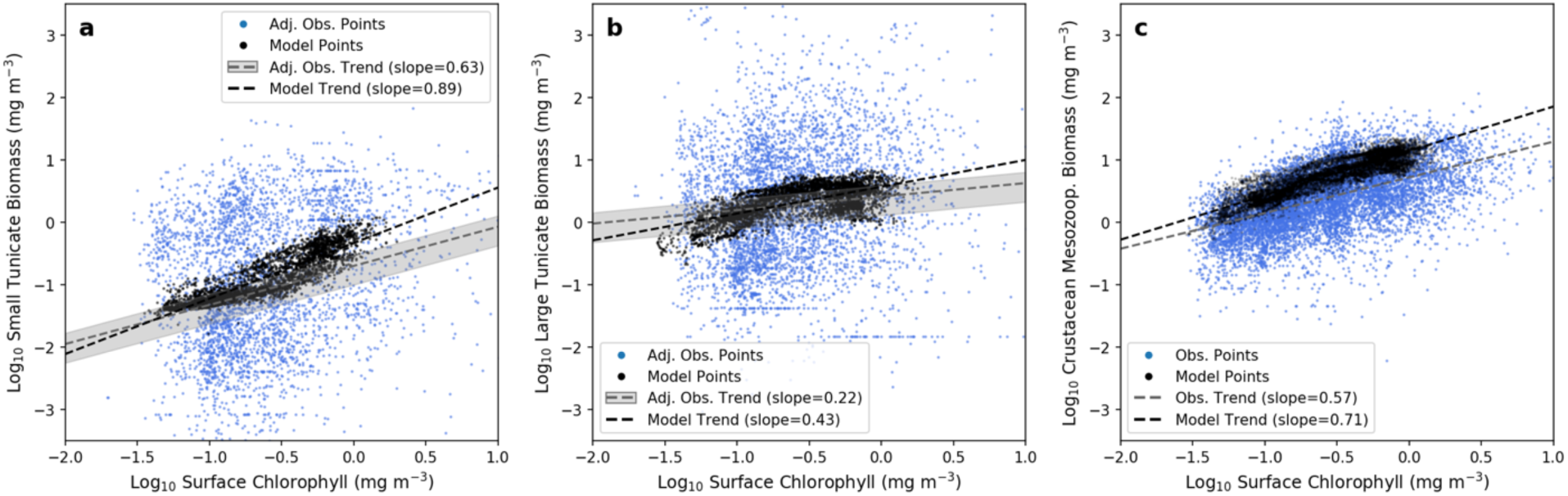
Log-log relationship between tunicate biomass and surface chlorophyll from adjusted observations (blue) and model data (black), sampled at the same locations as the observational dataset, for (a) small tunicates, (b) large tunicates, and (c) crustacean mesozooplankton. The observations were adjusted from the data compilation to account for the systematic low sampling bias from nets (10-fold adjustment), with the grey bars around the observational regression line (calculated from linear least squares regression) showing the 5-15x adjustment range. Model values are from the top 100-m, and the crustacean mesozooplankton biomass was computed as large mesozooplankton + 0.5*small mesozooplankton. Observational surface chlorophyll, as well as model data, were time means from the growing season.

The sensitivity tests illustrated the impact of various aspects of the base GZ-COBALT parameter choices, as well as the distinct physiology of tunicates relative to crustacean mesozooplankton. For large tunicates, the reduced basal respiration rate, relative to the literature-based mean, in the base GZ-COBALT simulation was key for achieving a mean biomass consistent with observations (see case 1, Fig. S3a-c). Similarly, for small tunicates, the lower maximum ingestion rate relative to the literature-based values, was also essential for achieving a mean biomass consistent with observations, and a biomass-chlorophyll scaling slope that did not deviate too far from the observational constraints (case 2, Fig. S3d-f). In case 2, total tunicate production also doubled, despite modest increases in biomass, largely due to the role of the small tunicates (Table S1). The tunicate biomass-chlorophyll scaling slope was a result of many factors, including competition between tunicate size classes, as the relative increase in small tunicate biomass resulted in a shallower scaling slope for large tunicates (case 2, Fig. S3e).

Sensitivity cases 3-5 focused on model formulations distinct to pelagic tunicates relative to the original crustacean mesozooplankton in COBALTv2. In case 3, where the ingestion half-saturation constant (with associated adjustments to the maximum ingestion rate) was set to the same value as that of the crustaceans, the resultant mean biomass and biomass-chlorophyll scaling for both tunicates matched the observations more closely (Fig. S3g-i). This was an interesting result and could have been a tuning choice; however, doing so would have negated a key criterion (of the half-saturation constant, K_i_, being much greater than the prey concentration) in converting measurements of clearance rate to ingestion rate. Setting the assimilation efficiency to a constant value (case 4) resulted in small tunicates being closer to observations, but large tunicates dropping significantly in biomass, particularly in the low productivity areas (Fig. S3j-l). This suggests that the variable assimilation efficiency was one factor in allowing large tunicates to survive in the subtropical gyres. Finally, in the case where large tunicate aggregation mortality was removed, this resulted in large tunicate biomass greatly increasing in the high chlorophyll areas, with associated increases in the biomass scaling slope (Fig. S3m-o). See Table S1 for a summary of the major results from the sensitivity cases as compared with the base GZ-COBALT simulation.

### 3.2 Seasonal cycle

All zooplankton exhibited a stronger seasonal cycle in the high chlorophyll seasonally stratified (HCSS) biome compared to the high chlorophyll permanently stratified (HCPS) and low chlorophyll (LC) biomes, with the biomass peak shifting later in the summer as zooplankton size increases. GZ exhibited a late summer (August-September) peak for both small and large tunicates. The large tunicates were also unique amongst zooplankton in that their biomass in the HCSS biome did not exceed that of the HCPS biome (Fig. 7). Results from the sensitivity cases showed that this is largely due to the large tunicate aggregation mortality, or jelly-falls (case 5, Fig. S4k-o), which serves to strongly dampen blooms. Additionally, reductions in the ingestion half-saturation constant (and associated maximum ingestion rate; case 3, Fig. S4a-e) and the constant assimilation efficiency (case 4) also reduced the magnitude of the blooms. Additionally, in case 4, the small tunicates’ bloom timing was also shifted to be slightly earlier (Fig. S4f-j). In the base GZ-COBALT configuration, the biomass of the non-GZ zooplankton were overall reduced compared to the COBALTv2 control (−7%, -6%, and -7.2% for small, medium, and large zooplankton, respectively), with the biggest difference seen in the summer microzooplankton biomass in the HCPS biome (Fig. 7a). Other substantial differences included the overall biomass of large crustacean mesozooplankton in the HCPS biome (Fig. 7c).

**Figure 7.**
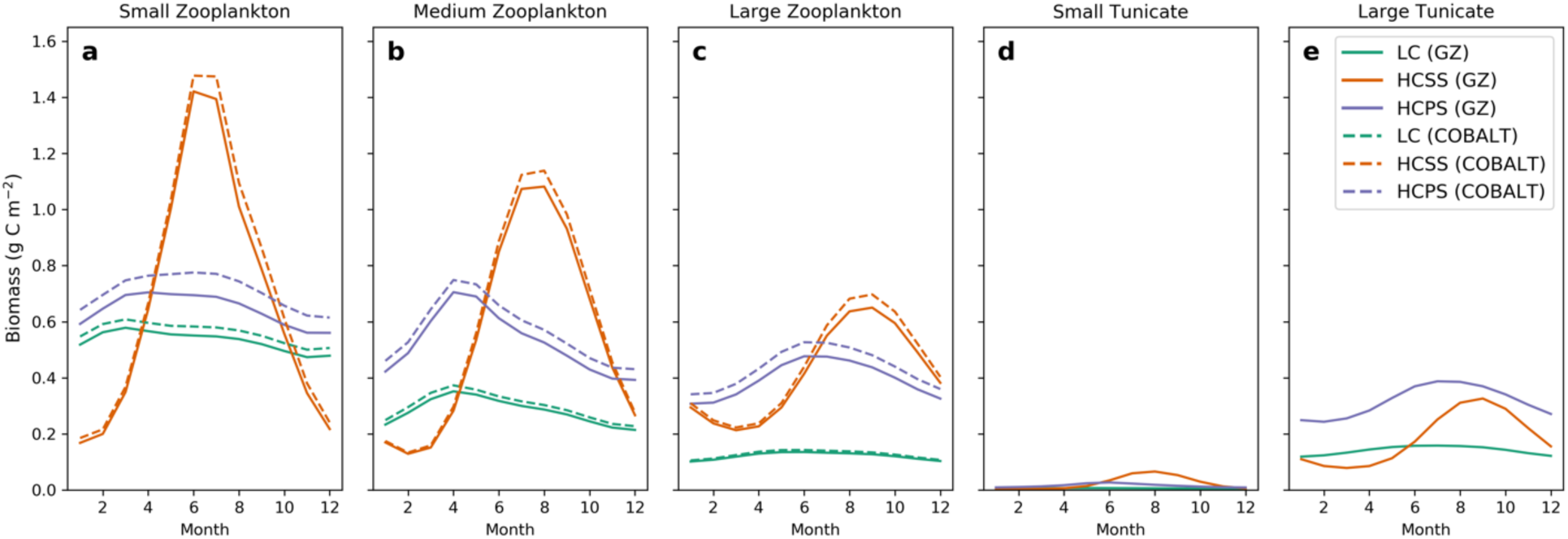
Seasonal cycle of modeled (a) microzooplankton, (b) medium zooplankton, (c) large zooplankton, (d) small tunicates, and (e) large tunicates, separated by biome. Biome definitions: LC = low chlorophyll, HCSS = high chlorophyll seasonally stratified, HCPS = high chlorophyll permanently stratified. Southern Hemisphere values were shifted six months such that Austral summer is represented by months 6-8. Solid lines indicate the GZ-COBALT simulation, and dashed lines show zooplankton values from the COBALTv2 control simulation.

### 3.3. Biogeochemical impacts

The overall impact of gelatinous zooplankton on the partitioning of energy between the microbial food web, export to depth, and energy available to higher trophic levels through mesozooplankton was assessed via the difference between GZ-COBALT and the control formulation (Fig. 8). This comparison suggests that the two tunicate classes have a competitive interaction with microzooplankton (Fig. 8c) and a small, but net negative impact of the total combined production of mesozooplankton (i.e., GZ and crustaceans, Fig. 8f). This is in spite of a competitive impact on crustacean zooplankton, which was greater for the small crustaceans, particularly in the upwelling zones, compared to the large crustaceans (Fig. S5).

**Figure 8.**
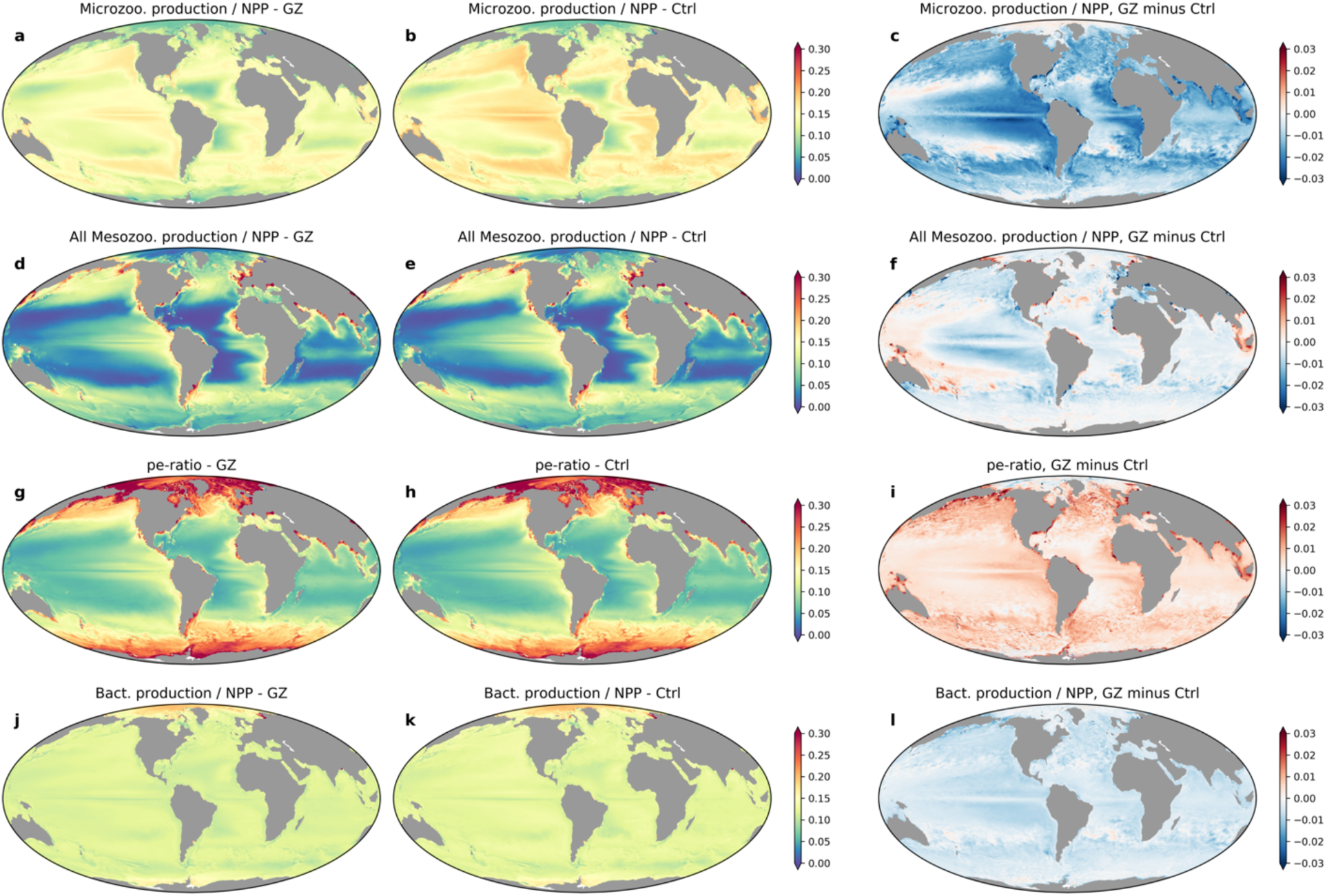
Differences in annual mean productivity ratios at the top 100 m between GZ-COBALT and the COBALTv2 control, showing ratios of microzooplankton production to NPP (a-c), all crustacean and tunicate mesozooplankton production to NPP (d-f), POC export past 100 m to NPP, or pe-ratio (g-i), and free-living heterotrophic bacteria production to NPP (j-l). The plots show the GZ-COBALT simulation (left column; a,d,g,j), the COBALTv2 control (center column; b,e,h,k), and the difference between the two (right column; c,f,i,l).

The differences between the simulations becomes more pronounced when considering plankton functional types that dominate recycling vs. export processes. With the addition of pelagic tunicates, the routing of carbon to the microbial food-web decreased, as indicated by declines in both the heterotrophic bacteria production ratio and the microzooplankton production ratio (Fig. 8c, 8l). Meanwhile, the particle export ratio (pe-ratio, defined as the export flux at 100 m divided by NPP) increased globally, averaging to a 5.3% increase (Fig. 8i). The total export flux at 100 m increased 2.1% from 6.23 Pg C y^-1^ to 6.36 Pg C y^-1^, but because of a corresponding slightly decline in NPP, the overall pe-ratio increase was greater (Table 4). This comes as small and large tunicates contributed 0.19 and 0.79 Pg C y^-1^, respectively, of total export production in the top 100 m, (8.88 Pg C y^-1^, Fig. 9), of which 72% sinks past 100 m. This increase in gelatinous-mediated export reflects a redistribution of export production from existing sources, with the largest coming from small mesozooplankton (Fig. 10a,c), as well as a reduction in the dissolved pool (8.2% decline in non-refractory dissolved organic carbon; Table 4).

**Figure 9.**
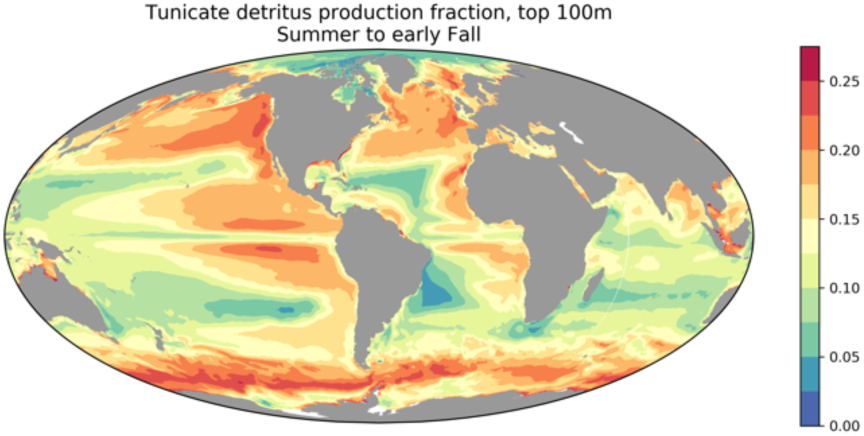
Fraction of detritus production in the top 100 m from tunicates in the summer and early fall months (June to September in the Northern Hemisphere, December to March in the Southern Hemisphere).

**Figure 10.**
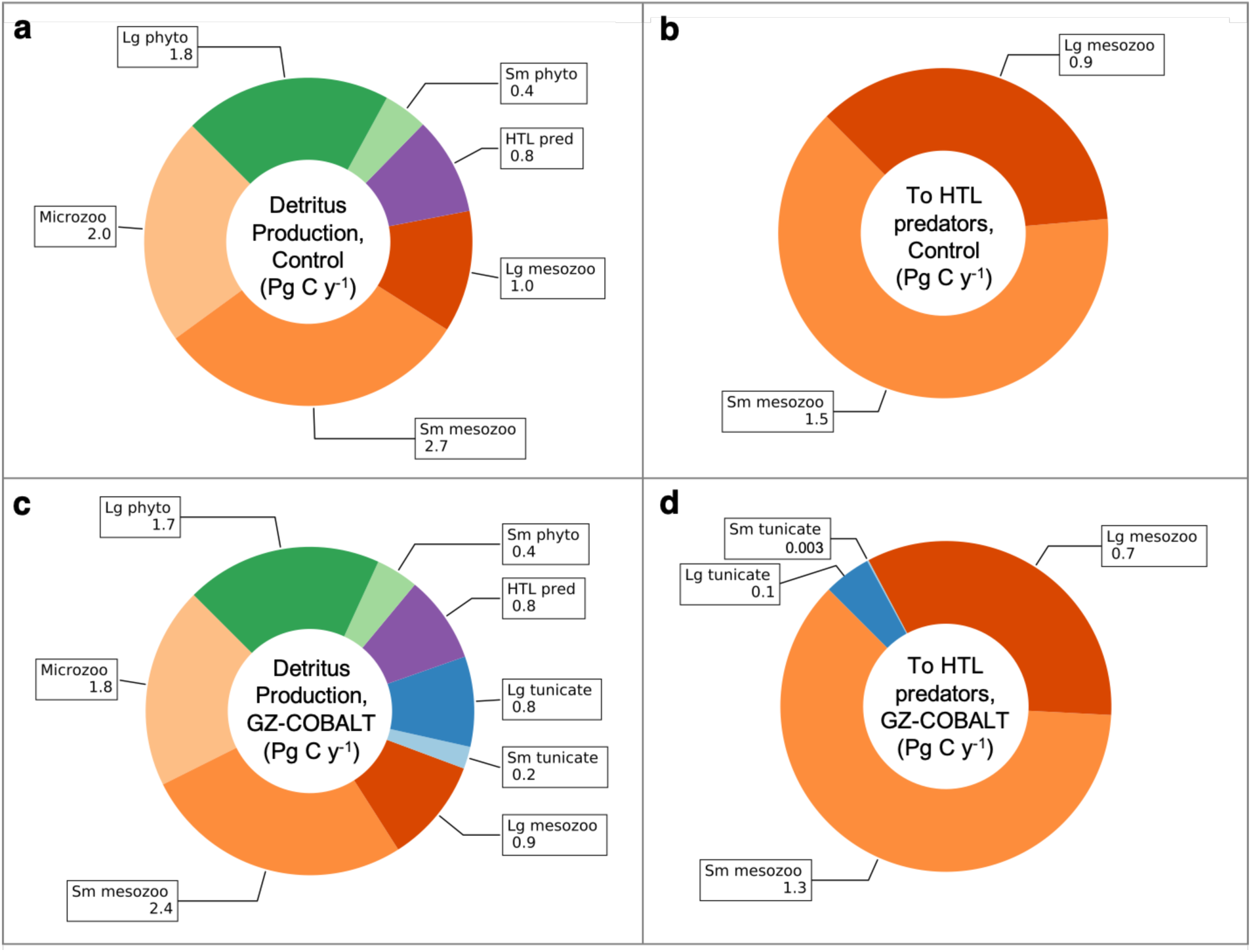
Top 100-m production of sinking detritus (a, c) and loss to higher trophic level (HTL) predators (b, d) in the COBALTv2 control (a, b) and GZ-COBALT (c, d) simulations. Of the total export production, approximately 72% of it sinks below 100 m. All values are from the top 100 m, and are in Pg C y^-1^.

## 4. Discussion

We have added a simple representation of two pelagic tunicate groups, representing appendicularians and thaliaceans, into the GFDL COBALTv2 ocean biogeochemistry model (GZ-COBALT) that captures large-scale patterns in tunicate distribution consistent with the emerging recognition of their importance to marine ecosystems, while maintaining a skillful representation of crustacean mesozooplankton, surface chlorophyll, and macronutrient concentrations. The GZ-COBALT simulation achieved a reasonable match between the modelled mean tunicate biomass and a global observational dataset, compiled from a range of sources including the COPEPOD database (O’Brien, 2014, 2005), Jellyfish Environmental Database Initiative (JeDI) (Condon et al., 2015), and KRILLBASE (Atkinson et al., 2017). Notably, GZ-COBALT captured a contrast between gelatinous and crustacean zooplankton types in their emergent relationship between biomass and surface chlorophyll (Fig. 6). These results confirm that it is possible to reconcile GZ biomass an order of magnitude above previous estimates (Remsen et al., 2004) with prevalent crustacean zooplankton populations: carbon flows through planktonic food-webs are sufficient to support both GZ and crustacean populations.

Observations of tunicate biomass exhibited high variability, even when compared with crustacean zooplankton observations gridded to the same horizontal resolution (Fig. 6) (Moriarty and O’Brien, 2013), which indicates either large sampling variability (e.g., from inconsistency in sampling effort and/or gear), unresolved physical or biological dynamics (Andersen, 1998; Boero et al., 2008; d’Ovidio et al., 2010; Greer et al., 2020; Lévy et al., 2018; Luo et al., 2014), or a combination thereof. Nearly all of the observations used in the present study were from net-based sampling systems, which destroy fragile gelatinous zooplankton. The comparison between nets and imaging systems from Remsen et al. (2004), while narrow in spatiotemporal extent, is one of the only robust comparisons that span the range of crustacean and gelatinous zooplankton, enabled in part by the larger sampling volume of their imaging system. Other studies have either focused on crustaceans and a few types of gelatinous zooplankton (Benfield et al., 1996; Stemmann et al., 2008) or other non-gelatinous plankton (Broughton and Lough, 2006; Cowen et al., 2013; Dennett et al., 2002). Benfield et al. (1996) found a 16-fold difference between appendicularian abundance as captured by imagery vs. nets, which was over five times greater of a difference than Remsen et al. (2004) found for appendicularians (3-fold difference in abundance). If anything, this indicates that the 10x adjustment for small tunicate carbon biomass we used may have been conservative. Broadly, our order-of-magnitude adjustment for tunicate biomass reflects our acknowledgement that this conversion is not precisely known. More high-quality observations of gelatinous zooplankton biomass are needed before biomass adjustments for net-based sampling can be more specific.

Nonetheless, the modelled variability for tunicates was lower than observations may suggest (cf. Stone and Steinberg, 2014), even when daily rather than monthly outputs were sampled (Fig. S6). This was also apparent for crustacean zooplankton, as it is a near-universal outcome when comparing global biogeochemical fields against point measurements or averages of small numbers of point measures (e.g., Krumhardt et al., 2017; Martiny et al., 2019; Saba et al., 2011; Usbeck et al., 2003). The discrepancy is admittedly acute for the tunicates, which was not unexpected given the sparsity and difficulty of measurements. A more complete understanding of the drivers of this patchiness and their implications will likely require high resolution physical simulations and GZ models capable of better resolving unique aspects of GZ life cycles and ecology conducive to patch formation (Groeneveld et al., 2020; Henschke et al., 2018a, 2018b).

In our analysis, we found a strong contrast in the biomass-chlorophyll relationship between crustacean zooplankton, small tunicates, and large tunicates, wherein the large tunicates exhibited a flatter scaling relationship compared to the steeper scalings of the small tunicates and crustaceans (Fig. 6). After incorporating an expanded view of GZ biomass considering undersampling by nets, the resultant observational biomass-chlorophyll scaling became one of our primary validation tools, as this emergent relationship can capture the mean biomass responses across productivity gradients. These relationships become important as steep spatial gradients in the contemporary ocean generally translates to amplified trends with climate change (Stock et al., 2017, 2014b). The shallower slope for large tunicates relative to crustacean zooplankton, in contrast, is indicative of less sensitivity to NPP, and suggestive of greater resilience to NPP declines projected by the majority of models under high emissions scenarios (Kwiatkowski et al., 2020) than their crustacean competitors. This would be consistent with current hypotheses for increased prevalence of GZ under climate change (Henschke et al., 2016).

### 4.1 Marine food web and biogeochemical impacts

Pelagic tunicates have long been identified as a potentially important source of carbon export, due to fecal pellets from salps (Iversen et al., 2017; Madin et al., 2006; Ramaswamy et al., 2005; Smith Jr et al., 2014; Urrère and Knauer, 1981) and appendicularians (Wilson et al., 2013), discarded appendicularian houses (Berline et al., 2011; Lombard and Kiørboe, 2010; Robison, 2005), and salp and pyrosome carcasses from jelly-falls (Henschke et al., 2013; Lebrato et al., 2013; Lebrato and Jones, 2009). Given the boom-and-bust population dynamic of pelagic tunicates, they can often be found to dominate POC export when present (Madin et al., 2006; Smith Jr et al., 2014). Indeed, a recent study from a NASA EXPORTS cruise found that salp fecal pellets comprised up to 80% of the detrital production in the upper 100 m in the NE Pacific when present, though they contributed an average of 28% of fecal pellet carbon production over a month-long sampling period (Stamieszkin et al., 2021). In our 20-year model climatology, large tunicate detritus production comprises 20% of the total detritus production in the top 100 m from summer to early fall in the same region (Fig. 9, S7). These values are a bit lower, but still fairly consistent with the sampled cruise mean, though the high observed variability in Stamieszkin et al. (2021) highlights the challenge in model-observation comparisons with snapshot studies at a single time point. Model comparisons with GZ-COBALT and sediment trap data, which integrates observations over longer time scales, will need to incorporate tunicate-specific POC sinking speeds and is a target for future work.

One common implication of observations of pelagic tunicate-mediated carbon export is that they would add to the existing POC export out of the surface ocean, often attributed to a combination of phyto-detritus and crustacean zooplankton fecal pellets (Buesseler et al., 2008; De La Rocha and Passow, 2007). Here, we found that when considering the upper oceans (top 100 m), the integration of pelagic tunicates with a “traditional” food-web model did not substantially increase total export flux past 100 m, which was 6.36 Pg C y^-1^ compared with 6.23 Pg C y^-1^ in the COBALTv2 control despite GZ accounting for 0.7 Pg C y^-1^. The integration of GZ thus led primarily to a redistribution of fluxes away from those previously attributed to crustacean zooplankton, rather than a creation of a new additive flux. The modest increase in particle export that did occur is consistent with compensation for reductions in dissolved organic carbon export arising from GZ-induced redirection of carbon flows away from the microbial food-web (Fig. 8, 9).

Compared to the offline estimates of tunicate export (1.3-3.9 Pg C y^-1^ at 100 m; Luo et al., 2020), the online GZ-COBALT model was lower, suggesting that food web and biogeochemical feedbacks decreased the overall export contribution of tunicates. Rather than relying on the direct application of GZ data, GZ-COBALT accomplishes this correspondence by mechanistically representing the primary observational features and satisfying myriad additional physical, biogeochemical, and food-web constraints. Large tunicates contributed about four times more export production than small tunicates in GZ-COBALT (0.79 vs. 0.19 Pg C y^-1^), with approximately 0.16 Pg C y^-1^ from jelly-falls (Table 4), which was only slightly lower than the offline estimates of 0.3-0.7 Pg C y^-1^. Tunicates in GZ-COBALT also contributed 0.1 Pg C y^-1^ to higher trophic level predators (Table 4, Fig. 10d), which was much lower than the offline estimates of 0.8-1.1 Pg C y^-1^ (Luo et al., 2020). The higher trophic level predation in the offline model was one of the least constrained parameters, as Luo et al. (2020) extracted a total GZ ecotrophic efficiency (fraction of production to predation) from a combination of EcoPath models (e.g., Ruzicka et al., 2020), and tuned this term for individual GZ groups to achieve a global fraction consistent with EcoPath estimates. Future observational and experimental work aimed to increasing our understanding of GZ predation by higher trophic levels should reduce the uncertainties associated with these global models.

The GZ-COBALT simulation showed that, compared with the COBALTv2 control, the largest impact of pelagic tunicates to ocean biogeochemical cycles is in the partitioning between the biological pump and the microbial food web. In GZ-COBALT, the impact of tunicates served to reduce microzooplankton and bacterial production as a function of NPP by 14% and 4%, respectively (Fig. 8). Pelagic tunicates, unlike other gelatinous zooplankton, are notable for primarily grazing on small particles and their high predator to prey size ratios (Conley et al., 2018; Sutherland et al., 2010), though some exceptions exist (Post, 2002; Walters et al., 2019). Recent work from Stukel et al. (2021) showed that in the Southern Ocean, the dominant salp, *S. thompsoni*, most strongly competed with protistan grazers instead of with krill due to the large size-based overlap between the salp and protistan diets. This is in contrast to previous speculation that salps are a dominant competitor of the Antarctic krill, *Euphausia superba*, and can be implicated as a factor in its long-term decline (Atkinson et al., 2004) and is consistent with recent evidence that this decline can be attributed to positive anomalies in the Southern Annular Mode (SAM) and loss of sea ice in the Southern Ocean (Atkinson et al., 2019). Our results indicate that while tunicates do compete with large crustacean mesozooplankton for prey, namely through the grazing of large phytoplankton and diatoms by appendicularians and doliolids, tunicates also serve as a source of food for both small and large crustacean mesozooplankton. Instead of competing with crustaceans, the magnitude of decline of microzooplankton and heterotrophic bacteria production in GZ-COBALT compared to the control and agreement with observations indicates that the microphagous tunicates serve as a trophic and carbon export shunt away from the microbial loop and towards the mesozooplankton food web and biological pump.

### 4.2 Model limitations

In this study, we focused on the effects of explicitly representing two groups of gelatinous zooplankton on the upper ocean ecosystem, the production of detritus, and how it relates to the balance between recycling and export in the euphotic zone. As such, we have simplified our model’s representation of detritus by grouping both slow and fast-sinking GZ detritus with the other modeled phyto- and zooplankton detritus, which all sink at a rate of 100 m d^-1^. Therefore, our estimates of the impact of modeled tunicates on export flux is conservative, compared to a model that explicitly represents fast-sinking tunicate detritus. We anticipate that explicitly representing fast-sinking large tunicate detritus could increase total POC export in GZ-COBALT by 0.1-0.2 Pg C y^-1^, which could further shift the balance between recycling and export. However, the net effect on upper ocean biogeochemical cycles and ecosystem functioning will likely not change substantially.

Amongst the marine zooplankton, thaliaceans are also notable for their complex life cycles which include the ability to reproduce asexually, alternation between sexual and asexual reproductive phases (salps and doliolids), and hermaphroditism (pyrosomes), all of which can yield large, transient, blooms under the right conditions (Andersen, 1998). Here, we have opted against modeling the complex life cycle of pelagic tunicates (Henschke et al., 2018a, 2015; Lombard et al., 2009b) for a more simple representation (Berline et al., 2011) aimed at capturing their mean state, seasonal fluctuations, and long-term trends that can be run in an Earth System Model for a few hundred years. As such, there were a number of necessary simplifications, and associated insights.

The model suggests that the mean turnover rate, as measured by the ratio of production over biomass, or P/B, for pelagic tunicates is overall lower (implying slower growth) than microzooplankton and crustacean mesozooplankton (Table 4, Fig. S8). While some shallow coastal areas exhibited P/B exceeding 0.1 d^-1^ in the summer for both small and large tunicates, the majority of the oceans had turnover rates < 0.03 d^-1^, even in the summer months. In contrast, the turnover rates for large tunicates as reported in the literature were 0.15-0.71 d^-1^ (Deibel, 1982; Gibson and Paffenhöfer, 2000; Madin and Purcell, 1992). While there may be some averaging due to the model’s monthly output, not even daily data captured the range of variability in the observations (Fig. S6). Future efforts may focus on determining whether the model’s inability to reproduce observed variability is due to its coarse horizontal resolution relative to the scales of observed variations in tunicate distributions (Greer et al., 2021; Luo et al., 2014), or due to the representation of the simplified life cycle. For some gelatinous zooplankton populations, a representation of the complex life cycle may be key for reproducing interannual and multi-decadal climate fluctuations (e.g., Henschke et al., 2018b).

### 4.3 Future outlook

Gelatinous zooplankton (GZ) are ubiquitous throughout the world’s oceans and a key contributor to marine food webs (Hays et al., 2018). Of the GZ, pelagic tunicates are likely the most important group in terms of carbon fluxes, due to their low trophic position and microphagous diet. We demonstrate, through a new model with food-web and biogeochemical feedbacks incorporated, that it is possible to reconcile an enhanced role of GZ in marine food webs with the established importance of crustacean mesozooplankton and other ocean biogeochemical constraints. Simulation results provide GZ flux estimates arising from a self-consistent physical-biological model satisfying myriad physical, biogeochemical and plankton food-web constraints, a unique contribution relative to previous “offline” estimates. Climate change is projected to drive decreases in NPP; coupled Model Intercomparison Project Phase 6 (CMIP6) models under the Shared Socioeconomic Pathway 5 (SSP5; fossil-fueled development) project a 3-9% decline in NPP by the year 2100 (Kwiatkowski et al., 2020). Associated with climate-induced NPP decreases, models also project a shift in mean pelagic body size: the abundance of large autotrophic phytoplankton will likely be reduced relative to their smaller counterparts due to increased warming, stratification, and subsequent nutrient limitation (Peter and Sommer, 2013). Consequently, as evidenced by the shallow scaling between biomass and chlorophyll, the role of large pelagic tunicates (thaliaceans) in marine food webs may further increase under climate change.

In this study, we have focused primarily on the upper ocean impacts of GZ, both to the food web and to the balance between recycling and export. Omitted in this work are considerations of the impact of fast sinking GZ export on the remineralization length scale and transfer efficiency to “sequestration depths”, which may have further impacts on benthic fluxes and air-sea CO_2_ exchange (Kwon et al., 2009; Lebrato et al., 2019; Luo et al., 2020; Sweetman et al., 2014; Titelman et al., 2006). In particular, there may be important feedbacks between climate-induced stratification and tunicate-mediated increases in export. Our results indicate that total carbon export was not significantly increased with inclusion of GZ in an ocean biogeochemical model (cf. Wright et al., 2021). However, these tunicate fluxes are globally quite significant and are associated with a redistribution of export from existing phytoplankton and mesozooplankton sources. As climate change will have differing impacts by taxonomic group, better understanding of the sources of carbon export and the mechanisms that drive their variation will improve our ability to project changes in the future.

## Acknowledgements

We thank colleagues who have contributed to the development and integration of various components of GFDL’s Earth System Model that make this work possible, as well as the technical support teams that maintain the NOAA/GFDL computing resources. JYL acknowledges support from the NOAA’s Marine Ecosystem Tipping Points Initiative. This work also benefitted from internal reviews by Elizabeth Drenkard and Cristina Schultz, and comments from two anonymous reviewers.

## Data availability

All model outputs necessary to reproduce the results in this manuscript, except the daily outputs, will be available upon publication at: https://doi.org/10.5281/zenodo.6533852. The daily outputs from the model needed to generate Fig. S6 are available upon request.

## Author contributions

JYL and CAS conceived and designed the study, and evaluated the model. JYL wrote the code, carried out simulations, analyzed data, and led the manuscript writing. JYL and TOB compiled data. All authors contributed critical feedback and edits to the final manuscript.

## Supplemental Material

### SI Text

#### Calculating biomass variance in the observational data

We followed the same procedure as Luo et al. (2020) in calculating the biomass variance in the observational data.

We calculated the geometric standard deviation for each biome. For a global estimate, since variance cannot be summed unless samples are completely independent, we first randomly sampled the biome-specific data using the lowest number of observations per biome as the sample number (appendicularians = 952, thaliaceans = 1121, crustacean mesozooplankton = 2392, all taxa = 952). The bulk geometric standard deviation was then calculated using this resampled dataset.

**Table S1.** Comparison of the major results from the GZ-COBALT base simulation vs. the sensitivity experiments. ‘z100’ refers to the top 100 m of the water column.

| Field | GZ-COBALT Base | GZ-COBALT Sensitivity Experiments |  |  |  |  |
| --- | --- | --- | --- | --- | --- | --- |
|  |  | Exp. 1 | Exp. 2 | Exp. 3 | Exp. 4 | Exp. 5 |
| NPP (Pg C y<sup>-1</sup>) | 53.7 | 55 | 51 | 54.5 | 54.5 | 53.6 |
| Detritus Production, z100 (Pg C y<sup>-1</sup>) | 8.88 | 8.71 | 8.97 | 8.81 | 8.81 | 8.91 |
| Export at 100 m (Pg C y<sup>-1</sup>) | 6.36 | 6.25 | 6.37 | 6.23 | 6.31 | 6.39 |
| Dissolved Organic Carbon (non-refractory), z100 (Pg C) | 7.41 | 7.91 | 6.33 | 7.68 | 7.79 | 7.24 |
| Total Grazing (Pg C y<sup>-1</sup>) | 38.8 | 39.7 | 37.5 | 39.3 | 39.3 | 38.8 |
| Zooplankton Ingestion, z100 (Pg C y<sup>-1</sup>) | 51.1 | 53.1 | 47.9 | 52.2 | 52.4 | 50.9 |
| HP Ingestion, z100 (Pg C y<sup>-1</sup>) | 2.18 | 2.33 | 1.96 | 2.26 | 2.25 | 2.27 |
| Total Phytoplankton Biomass (Pg C) | 0.482 | 0.486 | 0.466 | 0.485 | 0.486 | 0.48 |
| Total Zooplankton Biomass (Pg C) | 0.719 | 0.659 | 0.676 | 0.706 | 0.677 | 0.766 |
| Total Tunicate Biomass (Tg C) | 102 | 8.69 | 112 | 73 | 36.9 | 161 |
| Sm. Tunicate Biomass, z100 (Tg C) | 5.78 | 6.13 | 37 | 3.73 | 3.2 | 5.53 |
| Lg. Tunicate Biomass, z100 (Tg C) | 81.5 | 1.08 | 61.5 | 58.1 | 26.8 | 132 |
| Crustacean Mesozoop. (mdz + lgz) Biomass (Pg C) | 0.356 | 0.373 | 0.331 | 0.365 | 0.368 | 0.352 |
| Tunicate Prod., z100 (Pg C y<sup>-1</sup>) | 0.49 | 0.0361 | 1 | 0.259 | 0.0884 | 0.0724 |
| Crust. Mesozoo. (mdz + lgz) Prod., z100 (Pg C y<sup>-1</sup>) | 3.89 | 4.39 | 3.31 | 4.14 | 4.21 | 2.77 |
| Z-ratio | 0.072 | 0.08 | 0.065 | 0.076 | 0.077 | 0.07 |
| Sm. Tunicate Detritus Prod., z100 (Pg C y<sup>-1</sup>) | 0.19 | 0.21 | 1.58 | 0.11 | 0.14 | 0.18 |
| Lg. Tunicate Detritus Prod., z100 (Pg C y<sup>-1</sup>) | 0.79 | 0.01 | 0.51 | 0.5 | 0.38 | 0.96 |
| Tunicate Prod./Biomass, z100 ($y^{-1}$) | 5.61 | 5.02 | 10.2 | 4.19 | 2.95 | 5.25 |
| Crustacean Mesozoop. (mdz + lgz)<br>Prod./Biomass, z100 ($y^{-1}$) | 13.4 | 14.3 | 12.3 | 13.9 | 14 | 13.1 |
| Lg. Tunicate Prod. lost to HTL Pred. | 57% | 0% | 56% | 56% | 51% | 91% |
| Lg. Tunicate Prod. lost to Jelly-Falls | 35% | 4% | 33% | 34% | 34% | 0% |
| Slope of Sm. Tun/Chlorophyll scaling | 0.89 | 0.9 | 1.1 | 0.67 | 0.79 | 0.88 |
| Slope of Lg. Tun/Chlorophyll scaling | 0.43 | 0.26 | 0.29 | 0.39 | 0.62 | 0.58 |

**Figure S1.**
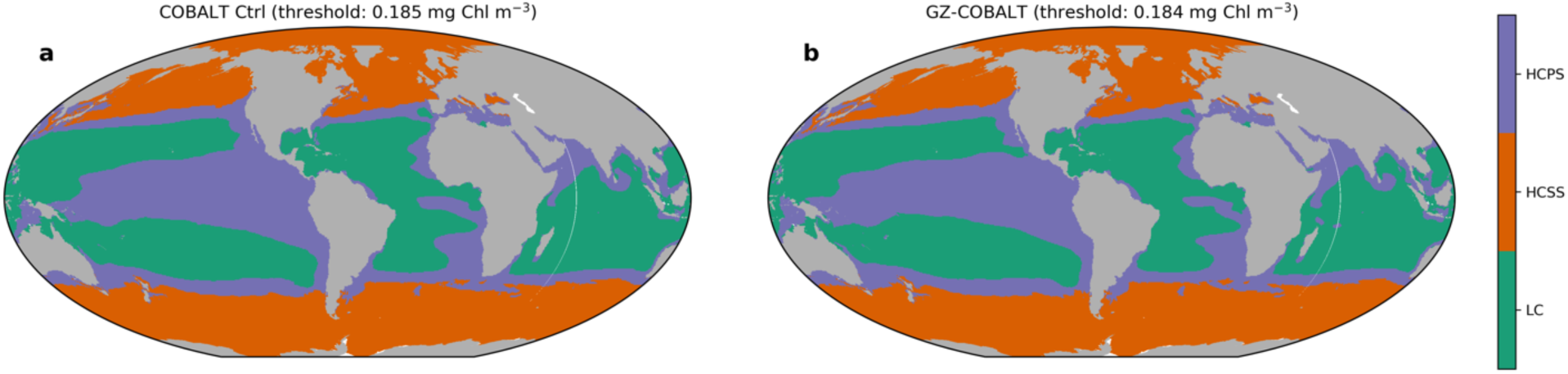
Major ocean biomes for the COBALTv2 control (a) and GZ-COBALT (b). The chlorophyll thresholds delineating the low chlorophyll (LC) regions were 0.185 and 0.184 mg Chl m^-3^ for the control and GZ-COBALT, respectively. The annual minimum of the mixed layer irradiance climatology (< 5 W m^-2^) delineated the high chlorophyll seasonally stratified (HCSS) biome from the high chlorophyll permanently stratified (HCPS) biome.

**Figure S2.**
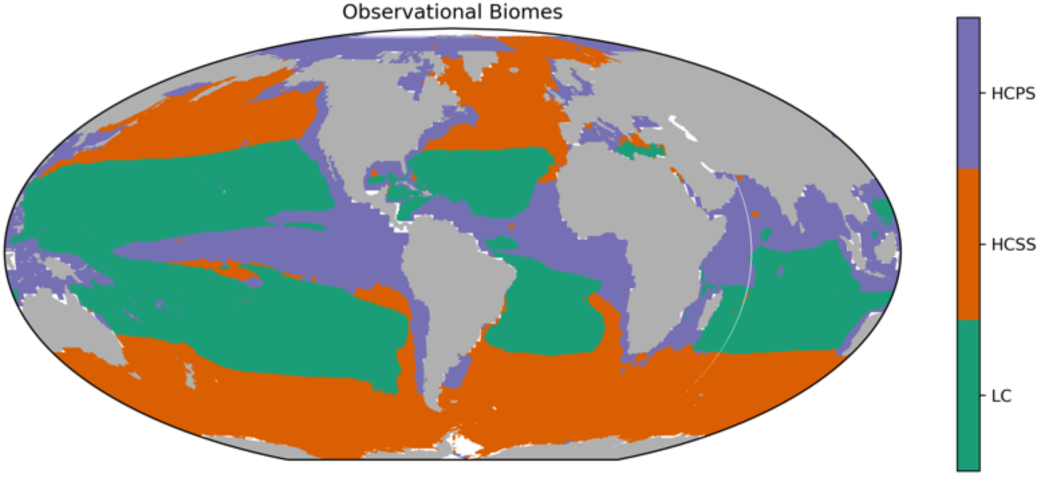
Major ocean biomes as mapped from observations. For observational chlorophyll, we used the GlobColour merged satellite chlorophyll product (MERIS, MODIS-Aqua, and SeaWiFS) monthly climatology for case 1 waters using the weighted averaging method, blended at latitudes south of 50°S with the Southern Ocean algorithm of Johnson et al. (2013). The threshold delineating the LC vs. the High Chlorophyll biomes was 0.125 mg Chl m^-3^. In contrast with the modelled biomes, mixed layer depth (MLD) was used to distinguish between HCPS and HCSS following Stock et al. (2014a). Observational MLD was from de Boyer Montégut et al. (2004) climatology, using the fixed density threshold criterion (0.03 kg m^-3^). The HCSS biome was marked by a maximum MLD from the annual climatology greater than 75 m.

**Figure S3.**
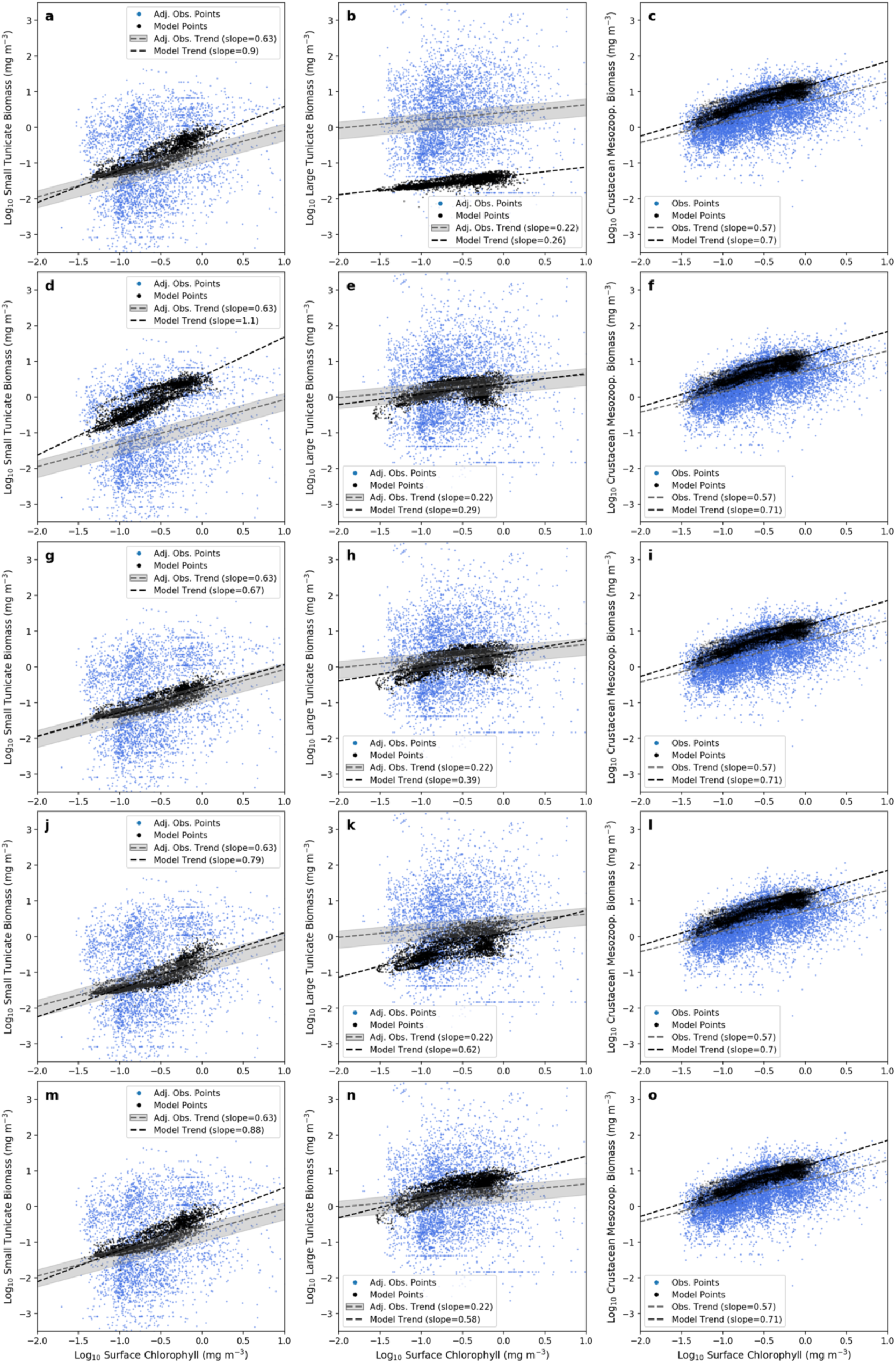
Biomass chlorophyll scaling for small tunicates (left column), large tunicates (center column) and crustacean mesozooplankton (right column) for the five sensitivity cases: case 1 (a-c), where large tunicate basal respiration is increased; case 2 (d-f), where small tunicate maximum ingestion rate is increased; case 3 (g-i), where the tunicate ingestion half-saturation constant is the same as crustaceans, and maximum ingestion is adjusted accordingly; case 4 (j-l), where tunicate assimilation efficiency is set to be a constant; and case 5 (m-o), where large tunicate aggregation mortality is turned off. See Table 2, and the caption on Fig. 6 for further details.

**Figure S4.**
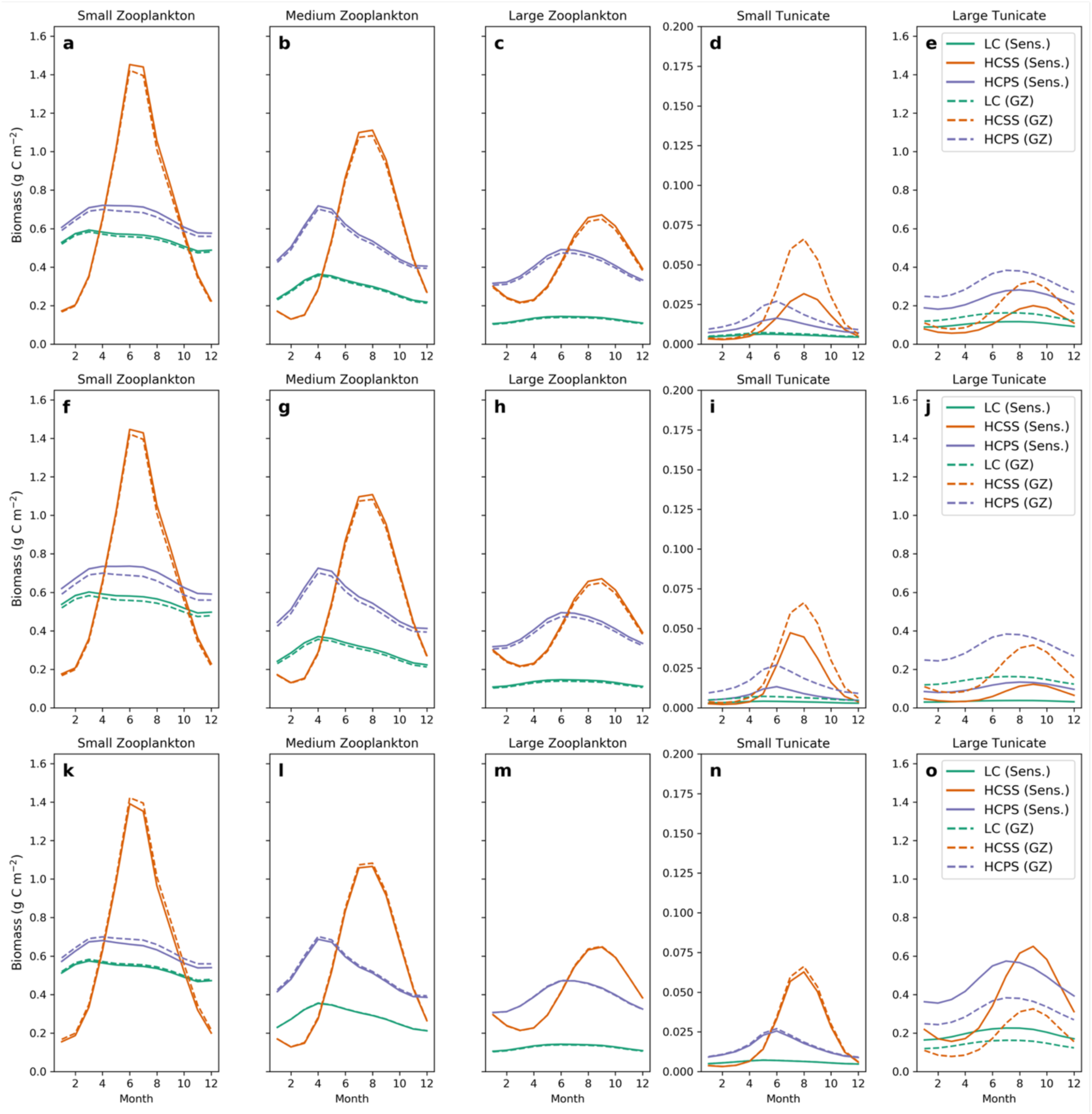
Zooplankton seasonal cycle differences between the GZ-COBALT base case (dashed) and three of the sensitivity cases: case 3 (a-e); case 4 (f-j); case 5 (k-o). See caption on Fig. 7 for further details.

**Figure S5.**
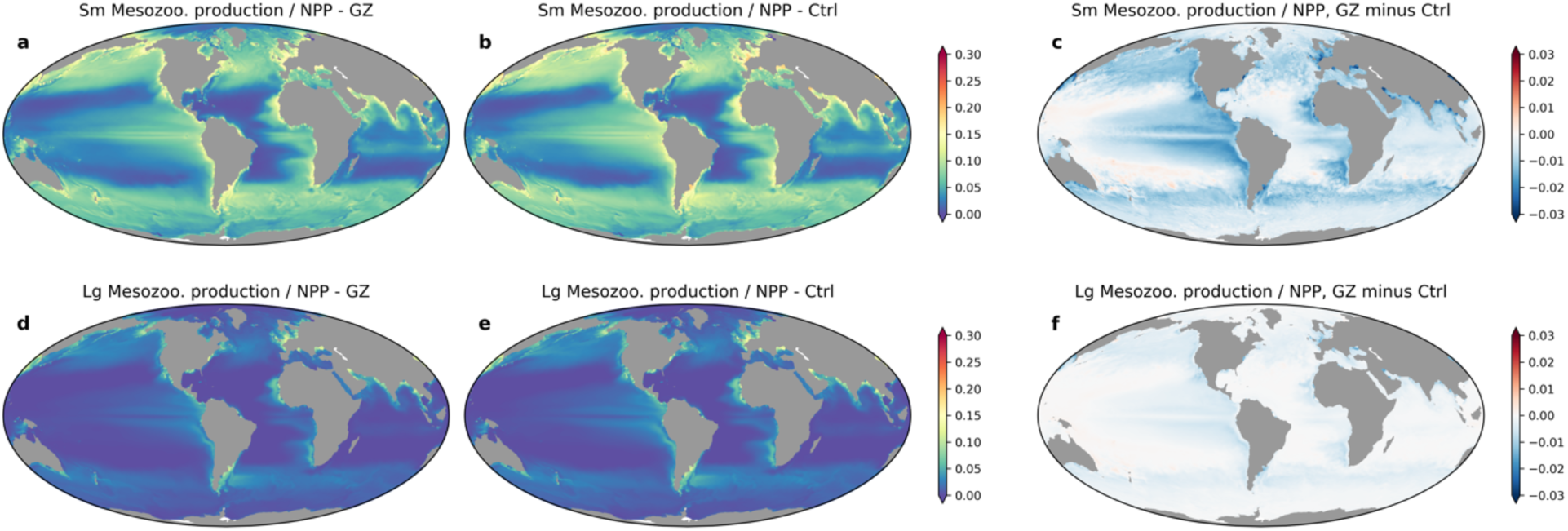
Differences in annual mean productivity ratios at the top 100 m between GZ-COBALT and the COBALTv2 control, showing ratios of small mesozooplankton (= medium zooplankton) production to NPP (a-c) and large mesozooplankton (= large zooplankton) production to NPP (d-f). The plots show the GZ-COBALT simulation (left column; a, d), the COBALTv2 control (center column; b, e), and the difference between the two (right column; c, f). Colorbar is set to be the same as in Fig. 8 to allow for a direct comparison.

**Figure S6.**
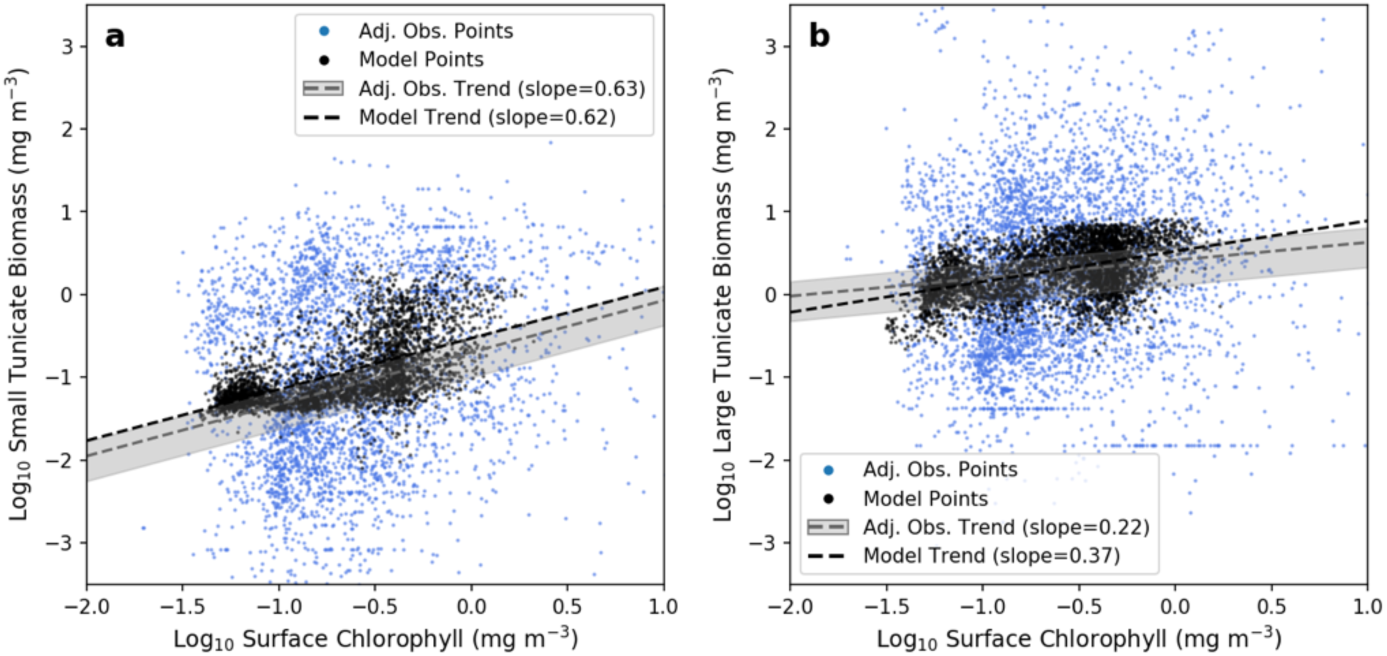
Biomass chlorophyll scaling for small (a) and large (b) tunicates, using daily output from the GZ-COBALT model, showing a randomly selected day within the growing season from the last year of the model simulation (2007). See also the caption in Fig. 6 for more details.

**Figure S7.**
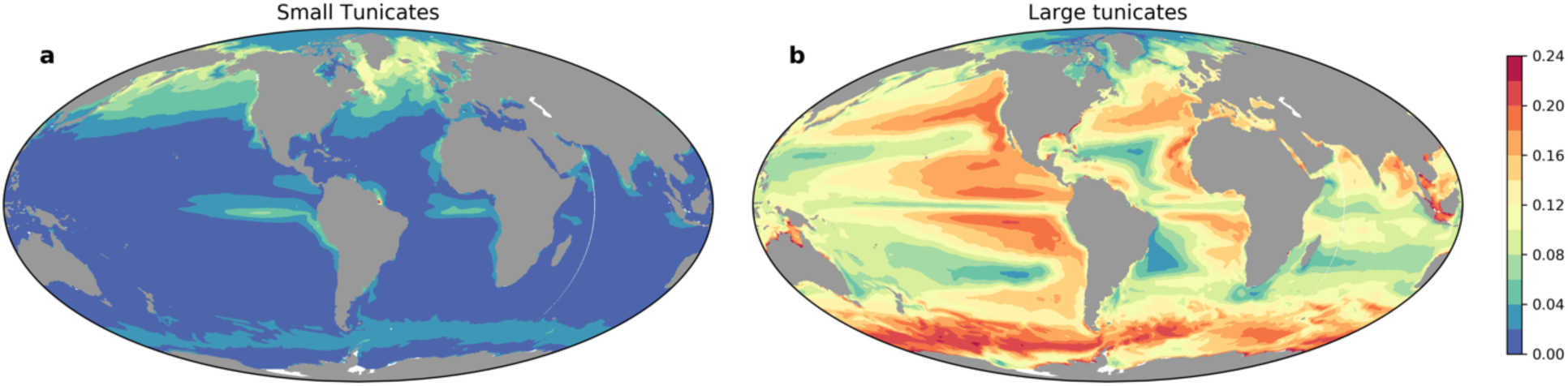
Fraction of summer to early fall detritus production in the top 100 m from (a) small tunicates and (b) large tunicates. Summer to early fall is defined as June-September in the Northern Hemisphere, and December-March in the Southern Hemisphere.

**Figure S8.**
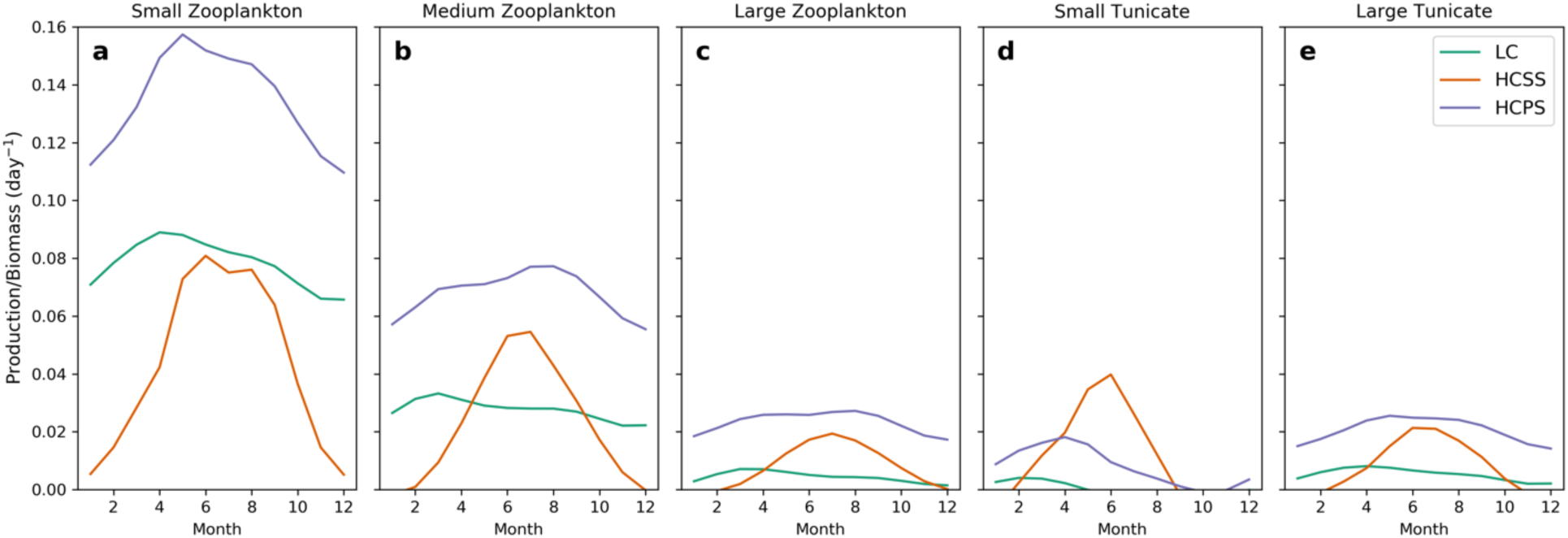
Mean daily Production/Biomass (P/B, d^-1^) ratios for small zooplankton (a), medium zooplankton (b), large zooplankton (c), small tunicates (d), and large tunicates (e). Months in the Southern Hemisphere are shifted such that Austral summer occurs during months 6-8, and Austral winter occurs during months 12, 1, and 2.

